# Miniaturized head-mounted device for whole cortex mesoscale imaging in freely behaving mice

**DOI:** 10.1101/2020.05.25.114892

**Authors:** Mathew L Rynes, Daniel Surinach, Samantha Linn, Michael Laroque, Vijay Rajendran, Judith Dominguez, Orestes Hadjistamolou, Zahra S Navabi, Leila Ghanbari, Gregory W Johnson, Mojtaba Nazari, Majid Mohajerani, Suhasa B Kodandaramaiah

## Abstract

The advent of genetically encoded calcium indicators, along with surgical preparations such as thinned skulls or refractive index matched skulls, have enabled mesoscale cortical activity imaging in head-fixed mice. Such imaging studies have revealed complex patterns of coordinated activity across the cortex during spontaneous behaviors, goal-directed behavior, locomotion, motor learning, and perceptual decision making. However, neural activity during unrestrained behavior significantly differs from neural activity in head-fixed animals. Whole-cortex imaging in freely behaving mice will enable the study of neural activity in a larger, more complex repertoire of behaviors not possible in head-fixed animals. Here we present the “Mesoscope,” a wide-field miniaturized, head-mounted fluorescence microscope compatible with transparent polymer skulls recently developed by our group. With a field of view of 8 mm x 10 mm and weighing less than 4 g, the Mesoscope can image most of the mouse dorsal cortex with resolution ranging from 39 to 56 µm. Stroboscopic illumination with blue and green LEDs allows for the measurement of both fluorescence changes due to calcium activity and reflectance signals to capture hemodynamic changes. We have used the Mesoscope to successfully record mesoscale calcium activity across the dorsal cortex during sensory-evoked stimuli, open field behaviors, and social interactions. Finally, combining the mesoscale imaging with electrophysiology enabled us to measure dynamics in extracellular glutamate release in the cortex during the transition from wakefulness to natural sleep.

## INTRODUCTION

Over the last several decades, neuroscientists have made significant strides to understand how neuronal computations occurring in isolated circuits in the brain mediate behavior. More recently, the advent of genetically encoded calcium^1^ and voltage indicators^2,3^, along with transgenic approaches for broadly expressing these indicators in the brain in a cell-type specific fashion^4,5^ has enabled mesoscale imaging of multiple cortical regions simultaneously^6–8^. These studies have revealed how neural activity across multiple regions of the cortex are coordinated in a variety of brain states and behaviors^9–15^.

Mesoscale cortical imaging studies have thus far been done exclusively in head-fixed animals^16,17^. Even relatively simple behavioral assays in awake head-fixed animals require significant time for acclimatization and training^18,19^. To overcome some limitations of head-fixation, immersive virtual reality environments^20^, voluntarily head-fixation of mice in their home cages^21^, and a moveable head-restraint apparatus^22^ have been used in mesoscale imaging studies. However, the lack of vestibular inputs and disruptions in eye-head movement coupling^23^,and behavioral effects from increased stress^24^ can significantly alter neural activity during head-fixation as compared to freely behaving animals^25^.

While several head-mounted, miniaturized imaging devices have been developed for imaging neural activity in freely moving animals^26–30^, these devices are designed to perform cellular resolution imaging, and thus have small field of view (FOV, < 1 mm^2^), limiting their application to imaging small brain regions. A head-mounted imaging device with a small FOV has recently been engineered for mesoscale imaging in rats^31^. In comparison to rats, mice have a much larger slate of genetic tools^2,32–34^. A head-mounted device with a large FOV for imaging the whole cortex in freely moving mice has the potential to significantly expand investigations of mesoscale cortical dynamics during free behavior.

Here, we introduce the ‘Mesoscope’, a miniature imaging device capable of simultaneously imaging multiple regions of the mouse dorsal cortex. The Mesoscope can image an 8 x 10 mm FOV with resolution ranging from 39.37 to 55.68 µm. The Mesoscope weighs 3.8 g and can be head-borne by a freely moving mouse. We have used the Mesoscope to successfully record mesoscale cortical activity in freely behaving solitary mice and during social interactions with a companion mouse.

## RESULTS

### Mesoscope Mechanical Design

The Mesoscope design was constrained by three main criteria. First, the overall weight of the device needed to be less than ∼15% of the mouse bodyweight (<4.0 g) to permit free behavior and mobility. Second, the device required the capability to image most of the dorsal cortex of the mouse. Third, we aimed to design a device with sufficient resolution to image mesoscale activity dynamics across the whole FOV. We recently developed See-Shells–transparent polymer skulls that can be chronically implanted on mice for very long (>300 days) durations of time and provide access to 45 mm^2^ of the dorsal cortex^35^. See-Shells were originally designed with screw holes to fasten a cemented titanium headpost for head-fixed imaging and protective cap. In this study, we adapted the See-Shell frame to fit the Mesoscope by incorporating a planar top surface, eliminating the headpost, and incorporating three tabs. Disk magnets were embedded in the two lateral tabs to align with disk magnets on the bottom of the Mesoscope (**Fig. 1a** bottom) or a protective cap, comprising a quick fitting mechanism. A short sleeve surrounding the Mesoscope base and the posterior tab in the bottom housing constrains the Mesoscope once mounted on the See-Shell. The tab at the back of the See-Shell frame gently restrains the mouse during removal of the protective cap and attachment of the Mesoscope. Attaching the Mesoscope typically takes less than 5 seconds and can be done without anesthetizing the mouse. For further stabilization, a fastening nut was embedded in the posterior tab of the See-Shell. An 0-80 screw is used to further secure the Mesoscope and minimize lateral motion.

**Figure 1:**
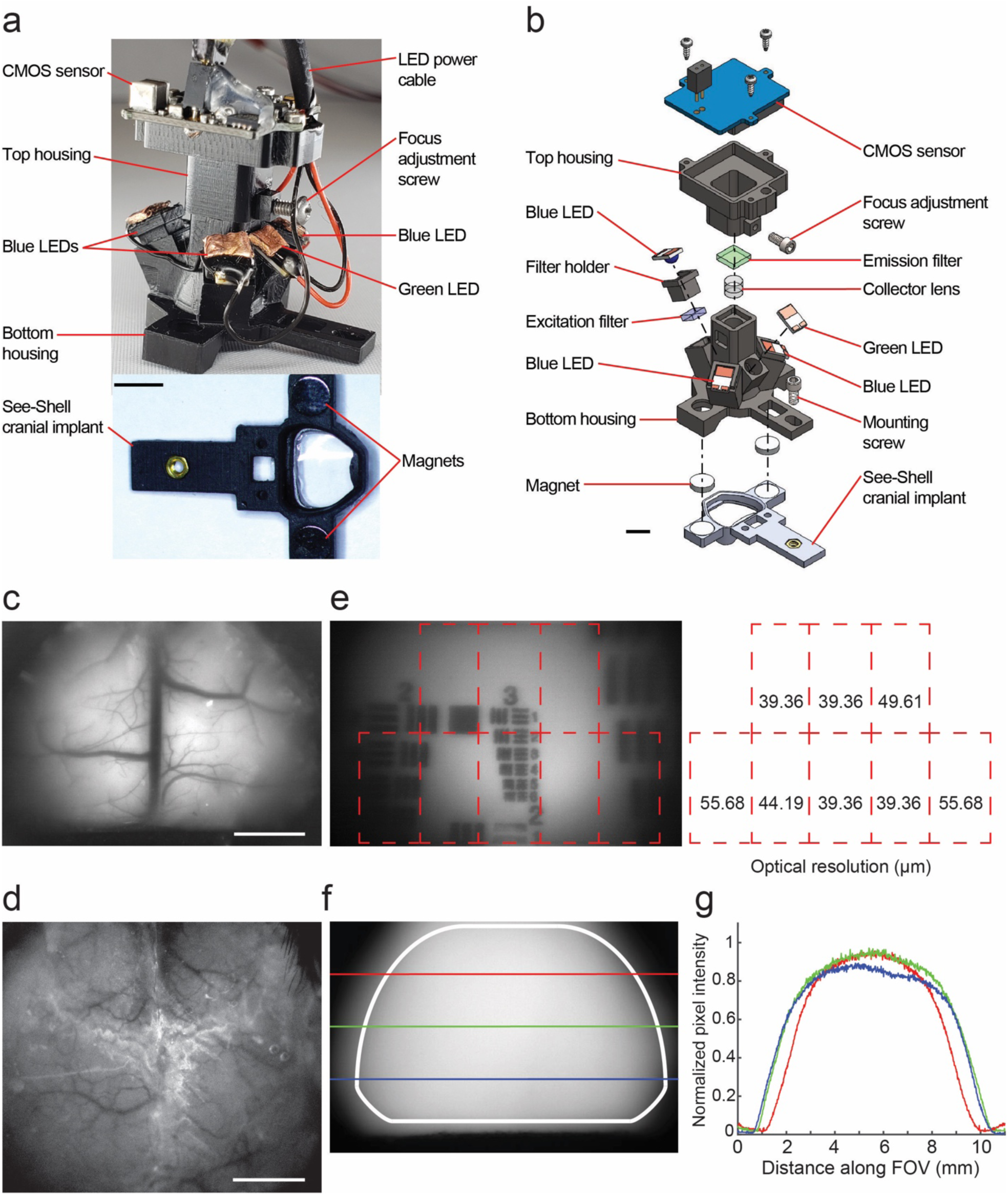
Mesoscope: A miniaturized head-mounted device for whole cortex mesoscale activity mapping in freely behaving mice: **(a)** Photograph of the fully assembled Mesoscope and the corresponding See-Shell implant. Scale bar, 5 mm **(b)** Computer-aided design (CAD) rendering of the Mesoscope showing the internal components. The device is attached to the See-Shell implant via magnets. Three blue LEDs are attached to a filter holder that has a 480 nm excitation filter to excite GCaMP6f in the cortex. The green LED is used to obtain reflectance measurements for hemodynamic correction. Resulting emission signals (∼520 nm) are focused through a collector lens and passed through an emission filter onto a CMOS sensor that can be manually focused. Scale bar indicates 5 mm. **(c)** Fluorescence image of the whole dorsal cortex of a Thy1-GCaMP6f mouse captured by the Mesoscope. Scale bar indicates 2 mm. **(d)** Comparative image of a head-fixed Thy1-GCcaMP6f mouse imaged using a standard epifluorescence macroscope through an intact skull preparation. Scale bar indicates 2 mm. **(e)** *Left:* Resolution test target overlaid onto the Mesoscope FOV. *Right:* Resolutions obtained within each specified grid. **(f)** Image of fluorescein dye infused agar phantom captured by the Mesoscope. Colored lines indicate medio-lateral sections along which illumination profiles were obtained FOV. **(g)** Plot of normalized illumination profiles from sections denoted in **f**.

The Mesoscope consists of two interlocking computer numerical control (CNC) machined Delrin housings (**Fig. 1a**). Three blue (480 nm) LEDs paired with an excitation filter (bandpass 470 nm) are installed into three illumination arms in the bottom housing (**Fig. 1b**). An additional green LED provides illumination for reflectance measurements. A biconvex collector lens and emission filter (bandpass 525 nm) are embedded in a central shaft of the bottom housing. A complementary metal oxide semiconductor (CMOS) imaging sensor is mounted on the top housing, which is designed to slide along the central square shaft of the bottom housing for focus adjustments. The CMOS sensor connects to a data acquisition (DAQ) board and is adapted from the open-source Miniscope system^26,27^. The three blue LEDs are wired in series and wires to power the green and three blue LEDs are routed through a commutator to alleviate torsional strain on the device when deployed on freely moving mice.

The overall weight of the device is 3.8 g, which is heavier than miniaturized microscopes with small FOVs^26,27^, but comparable to devices developed for volumetric imaging^29^. Based on computer-aided design (CAD) simulations, we estimate that the center of gravity is ∼24.7 mm above the mouse head. While shorter focal length optics could be used to reduce the length of the device, given the curvature of the cortex, lenses with shorter focal lengths result in optical distortion. Detailed instructions for assembling the Mesoscope are described in **Supplementary Figure 1** and **Supplementary Note 1**. CAD files for component fabrication and a bill of materials are provided in **Supplementary File 1**.

### Mesoscope Optical Performance

Typically, mesoscale cortical imaging is performed through an intact skull reinforced with refractive index matching transparent cement and a coverslip^36^. The optical resolution is dependent on the optical properties of the skull, the cement, and the quality of surgical preparation. In contrast, the Mesoscope images the cortex through a transparent polymer cranial window, so imaging resolution primarily depends on the collector lens within the Mesoscope. Qualitatively, imaging the cortex of a Thy1-GCaMP6f mouse^34^ through a transparent polymer skull with the Mesoscope (**Fig. 1c**) allowed us to achieve a high optical resolution across the FOV compared to imaging the cortex through the intact skull using an epifluorescence macroscope (**Fig. 1d**). This is demonstrated by the clearly visible vasculature at the surface of the brain in **Figure 1c**. To measure the resolution, we imaged a 1951 USAF resolution test target positioned at 8 different locations across the FOV. A representative image captured with group 3 of the test target located ∼ -5.5 mm anteroposterior (AP), ∼0 mm mediolateral (ML) is shown in **Figure 1e**. Lines in group 3, element 6 are clearly visible, indicating a resolution of 39.36 µm at this location. Since a single biconvex lens was used to image a convex surface, not all areas of the cortex are in focus and the optical resolution varied. In all experiments, the top housing was adjusted to obtain the best focus at ∼1 mm lateral to the midline. At this focus setting, the resolution ranged from 39 µm along the midline to 55.6 µm more laterally. Since our goal was to observe mesoscale calcium activity, we found these resolutions to be sufficient.

The Mesoscope’s array of three blue LEDs paired with excitation filters delivered a cumulative ∼31 mW of power to the brain. We imaged fluorescein dye-infused agar gel using the Mesoscope to investigate illumination uniformity (**Fig. 1f**). During initial testing, several versions of the bottom housing of the Mesoscope were rapidly prototyped using a stereo-lithography 3D printer. We iteratively changed the number of LEDs, their relative positions with respect to the optical axis, and their distance from the cortical surface. The final configuration consists of three blue LEDs slotted into square illumination arms. Two of the blue LEDs are oriented at 30 degrees with respect to the optical axis. The third, located at the anterior of the bottom housing, is oriented at 25 degrees with respect to the optical axis. The LEDs are revolved around the optical axis at angles of 90, 225, and 315 degrees. To reduce glare while maximizing power delivery to the FOV, the blue LEDs are 9.55 mm from the imaged surface. We observed that the normalized light intensity decreased by 56.1%, 53.3%, and 46.2%, compared to the maximum intensity, at the ML lines at 1.4 mm AP, -2.8 mm AP, and -4.2 mm AP, respectively (**Fig. 1g**). The greatest reduction of illumination from the maximum did not exceed 60%. These metrics are comparable to the performance of previously developed large FOV scopes^31^, and allowed signals obtained from all pixels to be well within the dynamic range of the CMOS sensor. A green LED was used to investigate reflectance for hemodynamic measurements^37^. The green LED delivered ∼0.22 mW of power and is located at the posterior of the bottom housing at an orientation of 55 degrees with respect to the optical axis.

The CMOS sensor captures images of the cortex alternatively illuminated by the blue LEDs filtered with a 20 nm band gap centered at 470 nm and the green LED. A trigger circuit uses time stamps of CMOS frame acquisitions to precisely switch between the blue LEDs and green LED (**Supplementary Fig. 2**). We found that the circuit and LEDs have first order dynamics. The blue LEDs and green LED have time constants (to reach 66.66% peak power) of 1.79 ± 0.31 ms and 1.97 ± 0.10 ms, respectively. The blue LEDs were powered on only during odd-numbered frames captured by the CMOS sensor. This was controlled by limiting switching on of the blue LEDs and green LED to 20 ms and 4 ms, respectively, starting after the CMOS sensor initiated each frame capture. It was also noted that the LEDs’ power intensity had a slow drift in average intensity value lasting ∼2 minutes after which they equilibrated (**Supplementary Fig. 2**). Therefore, we allocated a two-minute warm-up period for the blue LEDs during each experiment for the power intensity to stabilize before data collection.

### Comparison of Mesoscope to conventional wide-field epifluorescence macroscope

To benchmark *in vivo* imaging capabilities of the Mesoscope, we performed experiments in which a Thy1-GCaMP6f mouse anesthetized with isoflurane was imaged using the Mesoscope and a conventional epifluorescence macroscope in the same experimental session (**Fig. 2a**). Despite significant differences in optics quality, we captured qualitatively comparable images with both instruments. Vasculature was sharper in imaging data from the macroscope, but most major blood vessels and some finer details were visible in imaging data from the Mesoscope (**Fig. 2a**). Additionally, we observed heart beat-related oscillations in the calcium signals at 6-7 Hz in the imaging performed with both instruments (**Fig. 2b** and **c**), consistent with previous studies^31^. A discrete Fourier transform (DFT) of the calcium traces illustrates clear peaks that are observed in this frequency range (**Fig. 2c**). We next analyzed the DF/F traces captured from randomly chosen regions of interest (ROIs) at ∼ 3.5 mm laterally and ∼2 mm posterior to Bregma across the FOV. Qualitatively, the histograms of DF/Fs and z-score of intensity values acquired from three ROIs show minimal differences between the data captured from the two instruments (**Fig. 2d** and **e**).

**Figure 2:**
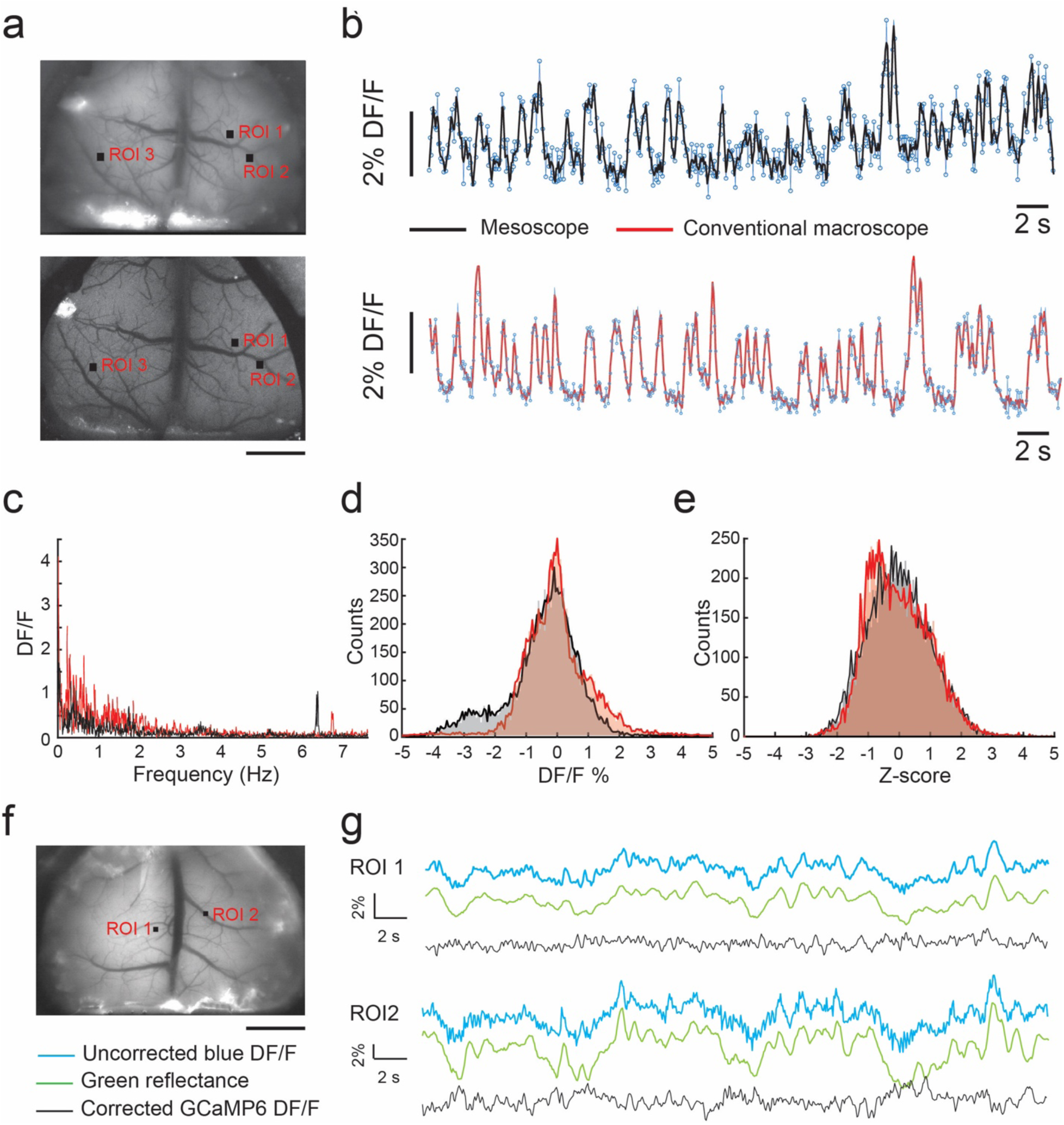
Comparison of calcium dynamics imaged with the Mesoscope to conventional widefield epifluorescence macroscope: **(a)** *Top:* Image of the dorsal cortex taken with Mesoscope of a Thy1-GCaMP6f mouse. *Bottom:* image taken with a conventional widefield epifluorescence macroscope during the same experimental session. Black squares indicate ROIs selected for DF/F traces analyzed in **b-e**. Scale bar indicates 2 mm. **(b)** DF/F traces of ROI 1 shown in **a** recorded for 120 seconds. Black and red traces indicate filtered traces (Savitzky-Golay filter, 3^rd^ order polynomial, 5 point moving average) of DF/F recorded from the Mesoscope and the macroscope respectively. Blue trace with markers indicates raw DF/F values under 1% isoflurane anesthesia. **(c)** Plot of the DF/F/Frequency obtained by computing the discrete Fourier series of the traces shown in **b. (d)** Histogram of DF/F values over 2-minute videos captured at the same 1% isoflurane anesthesia conditions with the Mesoscope and the conventional macroscope in the three ROIs indicated in **a. (e)** Histogram of z-scores of fluorescent intensity values over 2-minute videos captured at the same 1% isoflurane anesthesia conditions with the Mesoscope and the conventional macroscope in the three ROIs indicated in **a. (f)** Image of the dorsal cortex taken with Mesoscope of a Thy1-GCaMP6f mouse. Black squares indicate ROIs selected for hemodynamic corrections illustrated in **g**. Scale bar indicates 2 mm. **(g)** Uncorrected DF/F, reflectance DF/F and the hemodynamic corrected DF/F traces for the two ROIs shown in **f**.

Mesoscale imaging of calcium activity using GCaMP6f indicators is affected by the absorption of light by hemoglobin in the blood^37,38^, which can in turn significantly affect the measured fluorescence signal. Hemodynamic changes can be corrected using reflectance imaging. The green LED in the Mesoscope was switched on every alternate frame to capture reflectance changes simultaneously with calcium activity. **Figure 2g** shows representative traces of raw uncorrected blue DF/F signal and green reflectance DF/F signal from two randomly selected ROIs (**Fig. 2f**) in a freely behaving mouse as well as a ratiometric hemodynamic correction using a previously proposed methodology^37^. These results demonstrate that the Mesoscope can acquire calcium signals that are comparable to a conventional epifluorescence macroscope, and that the signal captured can further be corrected for artifacts due to hemodynamic response.

### Open field behavior of mice bearing head-mounted Mesoscope

We first assessed whether the implantation of the See-Shell and head-mounting the Mesoscope affected open field behavior. Mice bearing the Mesoscope were allowed to freely move in an open field arena for up to 5 minutes. Mice exhibited a repertoire of behaviors, including grooming and rearing, indicating their comfort with the Mesoscope (**Fig. 3a, Supplementary Video 1**). We performed behavioral tracking using an overhead camera in the arena, and then used traces from the recordings to quantify the mean speed, total distance, maximum speed and time spent in the center of the open field (**Fig. 3b-f**). Mice bearing the Mesoscope moved with an average speed of 4.56 ± 2.07 cm/s. In comparison, See-Shell implanted mice without the Mesoscope moved at an average speed of 4.94 ± 1.63 cm/s. There was no significant difference in average speed between both groups compared to the non-surgical control mice group, which moved with an average speed of 4.22 ± 1.24 cm/s (α = 0.05 for all tests, p = 0.569 between implant and control, p = 0.472 between Mesoscope and control, Student’s t-test). Similarly, we tracked the total distance moved in 5-minute trials. Mice bearing the Mesoscope moved a total distance of 1374.15 ± 623.40 cm. In comparison, See-Shell implanted mice without the Mesoscope moved 1492.65 ± 496.09 cm. There was no significant difference in total distance travelled between both groups compared to that from control mice (p = 0.059 between implant and control, p = 0.4937 between Mesoscope and control). Mice bearing the Mesoscope had an average maximum speed of 49.08 ± 16.48 cm/s. In comparison, See-Shell-implanted mice without the Mesoscope had an average maximum speed of 53.06 ± 8.75 cm/s. There was no significant difference in maximum speed between both groups and the control mice (p > 0.05 between implant and control, and between Mesoscope and control, Student’s t-test). There was a statistically significant difference between the time control mice spent in the open field versus mice with the See-Shell implant and mice with the Mesoscope (p = > 0.05 between Mesoscope and control, and between implant and control). This could be caused by the inability of the mice to investigate the corners of the arena with the Mesoscope or See-Shell implanted. Tests of how the Mesoscope or See-Shells affect anxiety levels would require the open field arena test to be used in conjunction with other behavioral tests^39^. These results indicate that the Mesoscope has little to no effect on the free behavior of the mice, consistent with results obtained with other miniaturized head-mounted imaging devices^29^.

**Figure 3:**
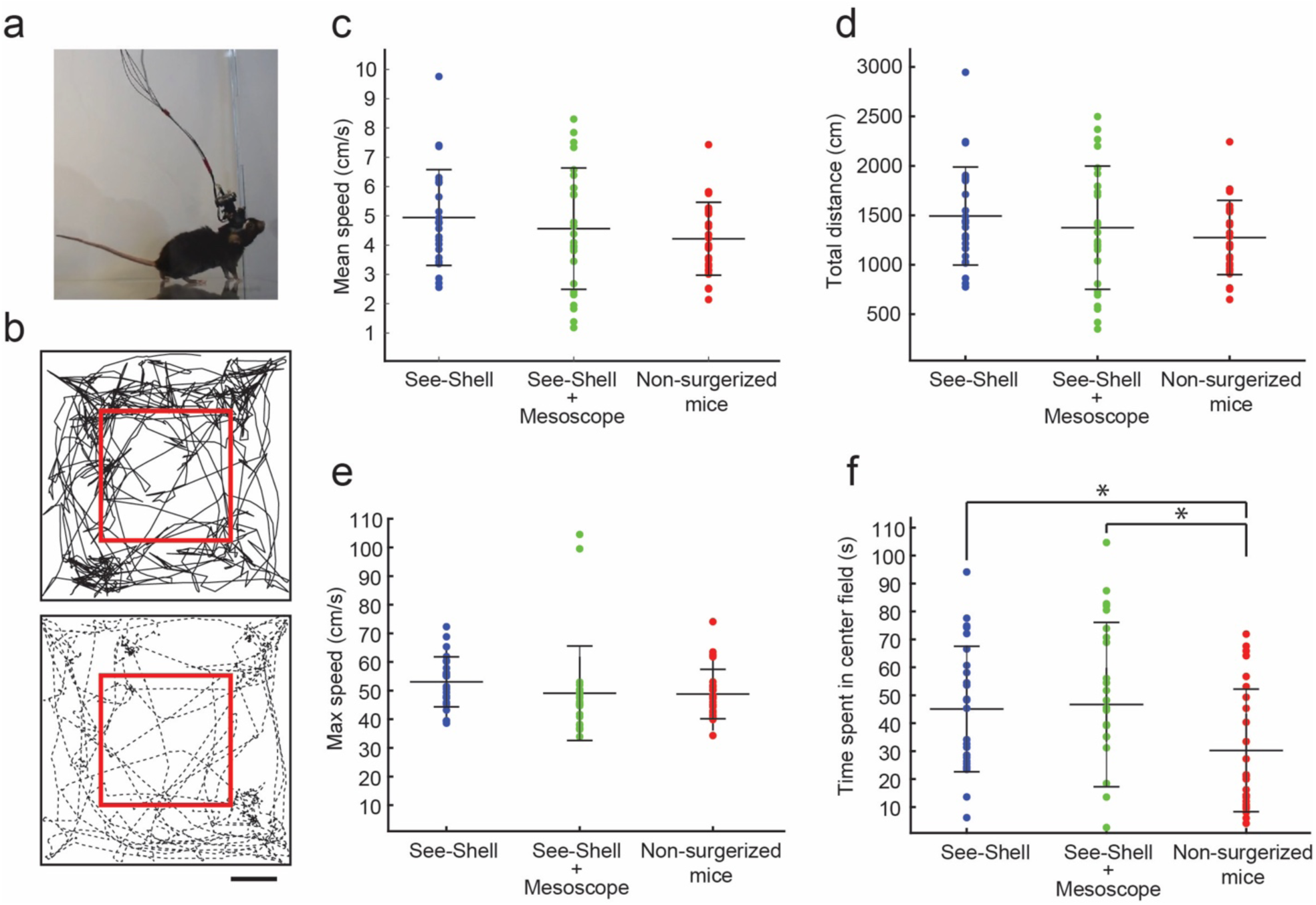
Head-mounting the Mesoscope has minimum effect on open field behavior: **(a)** Photograph of a mouse bearing a Mesoscope in an open field behavioral arena. Scale bar 10 cm. Mice exhibit a repertoire of complex behaviors with the Mesoscope attached (**Supplementary Video 1**). **(b)** Trace of mouse position in a 5-minute trial. *Top:* See-Shell implanted mouse without the Mesoscope. *Bottom:* Mouse bearing head-mounted Mesoscope. Red box indicates region of the arena denoted as the open field. Scale bar indicates 5 cm. **(c)** Plot of mean speed of mice with a See-Shell implanted, mice with both the See-Shell and head-mounted Mesoscope, and non-surgerized control mice. **(d)** Plot of total distance traveled by mice with a See-Shell implanted, mice with both the See-Shell and head-mounted Mesoscope, and non-surgerized control mice. **(e)** Plot of maximum speed of mice with a See-Shell implanted, mice with both the See-Shell and head-mounted Mesoscope, and non-surgerized control mice. **(f)** Plot of time spent at the center of the open field by mice with a See-Shell implanted, mice with both the See-Shell and head-mounted Mesoscope, and non-surgerized control mice. * indicates p < 0.05, t-test.

### Imaging sensory stimulus evoked responses across the cortex

Stimulating distinct sensory pathways evokes neural activity in distinct primary sensory areas located within the dorsal cortex. We used the Mesoscope to map evoked responses to somatosensory and visual stimuli. A mouse under light (< 1% isoflurane) anesthesia was first provided with a series of brief vibrational stimuli (1s long, 100 Hz) to the right hind limb (**Fig. 4a**). As expected, somatosensory stimulus evoked robust calcium activity in the contralateral hind limb (HL) region of the somatosensory cortex within 500 ms of the onset of stimulus (**Fig. 4b** and **c**). The peak post stimulus response was 1.68% ± 0.49% DF/F (**Fig. 4d**, n = 17 trials in 1 mouse). In comparison, the peak post-stimulus response on the ipsilateral side was 0.70% ± 0.34% DF/F (**Fig. 4e**), significantly lower as compared to the contralateral response (**Fig. 4f**, p < 0.01, t-test). In a similar fashion, the same mouse was presented with a 100 ms long flash of white light to the left eye, and we observed a robust increase in calcium activity in the contralateral visual cortex (**Fig. 4h** and **i**). The peak post stimulus response was 1.7% ± 0.32% DF/F (**Fig. 4j**, n = 18 trials). In comparison, the peak post-stimulus response on the ipsilateral visual cortex was 0.38% ± 0.057% DF/F (**Fig. 4k**), significantly lower as compared to the contralateral response (**Fig. 4l**, p < 0.01, t-test). Thus, the Mesoscope can reliably measure expected evoked response to varied sensory stimuli in both hemispheres of the dorsal cortex.

**Figure 4:**
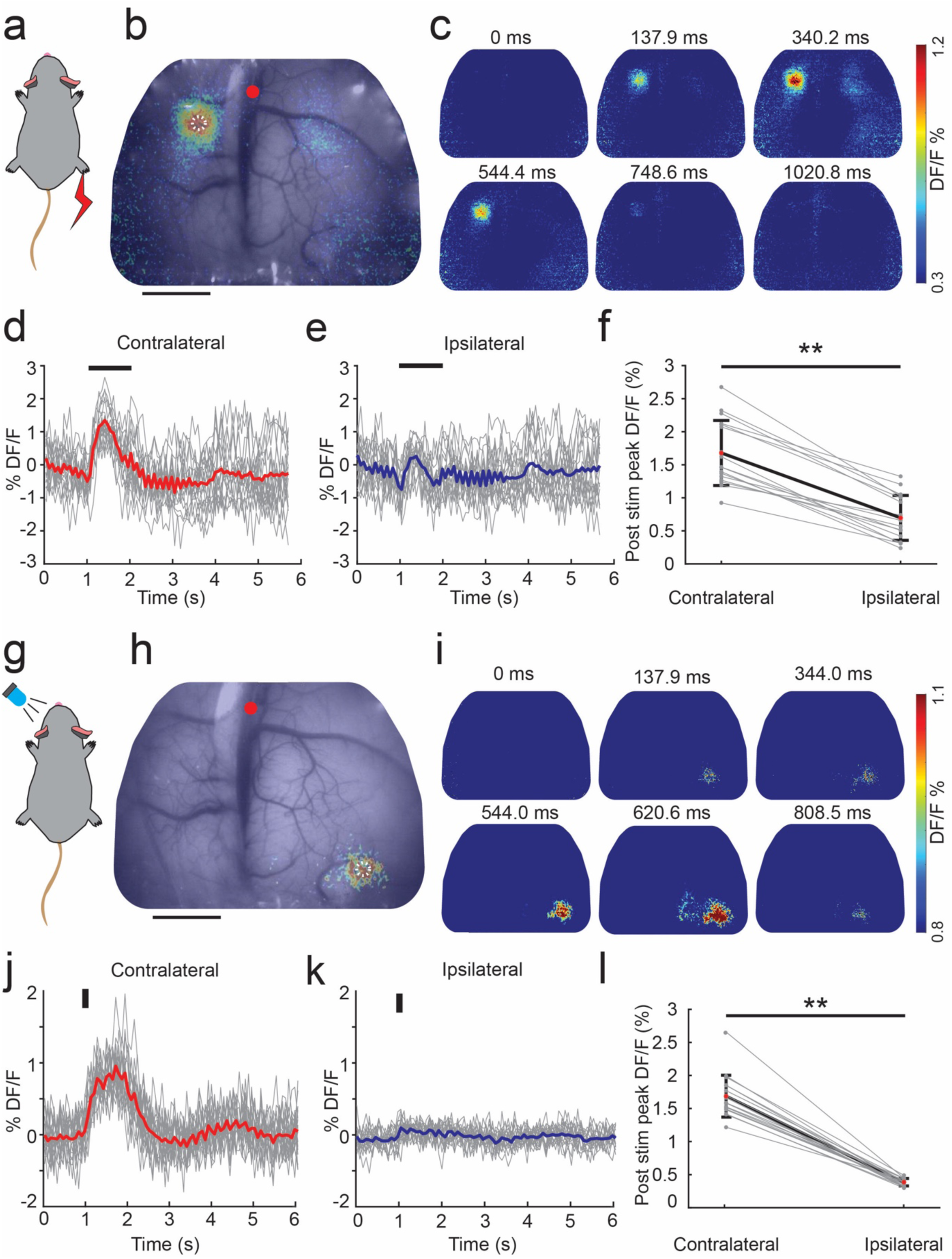
Sensory stimulus evoked responses imaged by the Mesoscope: **(a)** Depiction of a Thy1-GCaMP6f mouse under 1% isoflurane anesthesia conditions with a vibrational stimulus applied to the right hind limb. **(b)** Composite image of the raw, grayscale brain captured through the Mesoscope and the pseudo-color frame where the largest average DF/F occurred within the 1-second stimulus period. White circle indicates ROI analyzed in **d** and **e**. Red dot indicates Bregma. Scale bar 3 mm **(c)** Pseudo-color images of time-series evolution of the calcium response in the average DF/F video in response to the vibration stimulus. **(d)** DF/F traces of the contralateral ROI drawn in **b**. Red line denotes the average contralateral response whereas the grey lines denote each individual trial. Black bar shows the time and duration of the vibrational stimulus. **(e)** DF/F traces of ipsilateral pair of the ROI drawn in **b**. The blue line denotes the average ipsilateral response whereas the grey lines denote each individual trial. **(f)** Peak contralateral and ipsilateral response to the vibration stimulus taken within a 1-second window after the stimulus is applied. The bolded line corresponds to the average peak response whereas the grey lines indicate the peak response for each trial. ** indicates p < 0.01, t-test. **(g)** Depiction of the same mouse as in **a** to **f** with a visual stimulus applied to the left eye. **(h)** Composite image of the raw, grayscale brain captured through the Mesoscope and the pseudo-color frame where the largest average DF/F occurred after stimulus. White circle indicates ROI analyzed in **j** and **k**. Scale bar 3 mm. **(i)** Pseudo-color images of time-series evolution of the calcium response in the average DF/F video in response to the visual stimulus. **(j)** DF/F traces of the contralateral ROI drawn in **h**. Red line denotes the average contralateral response whereas the grey lines denote each individual trial. Black bar shows the time and duration of the visual stimulus. **(k)** DF/F traces of ipsilateral pair of the ROI drawn in **h**. The blue line denotes the average ipsilateral response whereas the grey lines denote each individual trial. **(l)** Peak contralateral and ipsilateral response to the visual stimulus taken within a 1-second window after the stimulus is applied. The bolded line corresponds to the average peak response whereas the grey lines indicate the peak response for each trial. ** indicates p < 0.01, t-test.

### Stability of Mesoscope imaging in freely moving mice

We next proceeded to use the Mesoscope to obtain calcium dynamics in freely behaving mice. Free behavior can cause motion artifacts due to two issues: the Mesoscope’s movement relative to the implant or the brain’s movement relative to the implant. When the Mesoscope was attached with the mounting screw on the rear tab, there were minimal lateral motion artifacts while the mouse was performing vigorous behaviors such as rearing and grooming (**Supplementary Video 2)**. In the trials that were analyzed, the absolute maximum x and y displacements of the FOV were 21.0 ± 21.8 µm and 14.4 ± 17.1 µm, respectively (**Supplementary Fig. 3**, n = 3 mice, 3 trials each). The mean absolute x and y displacement were 2.0 ± 2.5 µm and 9.3 ± 4.9 µm, respectively. These motion artifacts were adequately corrected for using a rigid body motion correction algorithm^40^.

Limitations on the weight of the device precluded us from adding any more instrumentation such as a laser range finder^21^ to measure z-axis deflection. However, based on the numerical aperture of the collector lens, the Mesoscope’s depth of focus ranges from ∼120 µm near the optical axis to ∼180µm in more lateral regions of the FOV. Thus, small fluctuations in the z-direction should be within the depth of focus of the device. While it is possible for the brain to move relative to the implant, our previous study has demonstrated that the brain is stable enough across much of the dorsal cortex to enable 2P imaging of single neurons in awake head-fixed animals^41^. Fluctuations of the brain position relative to the implant larger than 100 µm should be easily perceptible to the naked eye. In this study (**Supplementary Video 2**), as well as our previous work^41^, such large fluctuations have not been observed.

### Tracking changes in superior sagittal sinus diameter during free behavior

The Mesoscope images the cortex through the transparent See-Shell implants, allowing relatively high-resolution imaging of the surface cerebral vasculature compared to intact skull imaging. In raw calcium activity imaging videos acquired during freely moving behavior, we observed large changes in the diameter of the superior sagittal sinus (SSS) that were perceptible to the naked eye (**Supplementary Video 2**). Using image processing, we analyzed the changes in diameter of the SSS at various locations (**Fig. 5a**). Representative traces of the time-series of SSS diameter during an open field behavior trial are shown in **Figure 5b**. The measured diameters varied throughout the trial and the variance in percentage change in vessel diameter was comparable when the mice remained stationary, were moving or grooming (n = 6 trials, 3 mice) (**Fig. 5c**). There was no statistically significant difference between the groups (p > 0.05, Wilcoxon signed-rank test). The variance increased during social contact (n = 3 trials, 3 mice) (**Fig. 5d**). In all trials, we further observed rapid (<0.5 s) and large (>10%) contractions in the SSS. We initially hypothesized that they may occur during motion artifacts caused by behavior. But such spike-like reductions in diameters were observed even when the mice were stationary (**Fig. 5e**). Like the variances, there was no statistically significant difference between the spike frequencies (p > 0.05, paired sample t-test). A consequence of this observed phenomenon was that the areas adjoining the SSS had large lateral motion artifacts that could not be corrected using rigid body motion correction algorithms. Thus, areas within ∼500 µm were excluded in our data analyses of calcium activities. Further, blood from much of the frontal and parietal areas drain into the SSS. Fluctuations in the diameter of the SSS may affect the outflow of blood from these upstream brain regions and potentially points to a mechanism for mesoscale control of cerebral blood flow.

**Figure 5:**
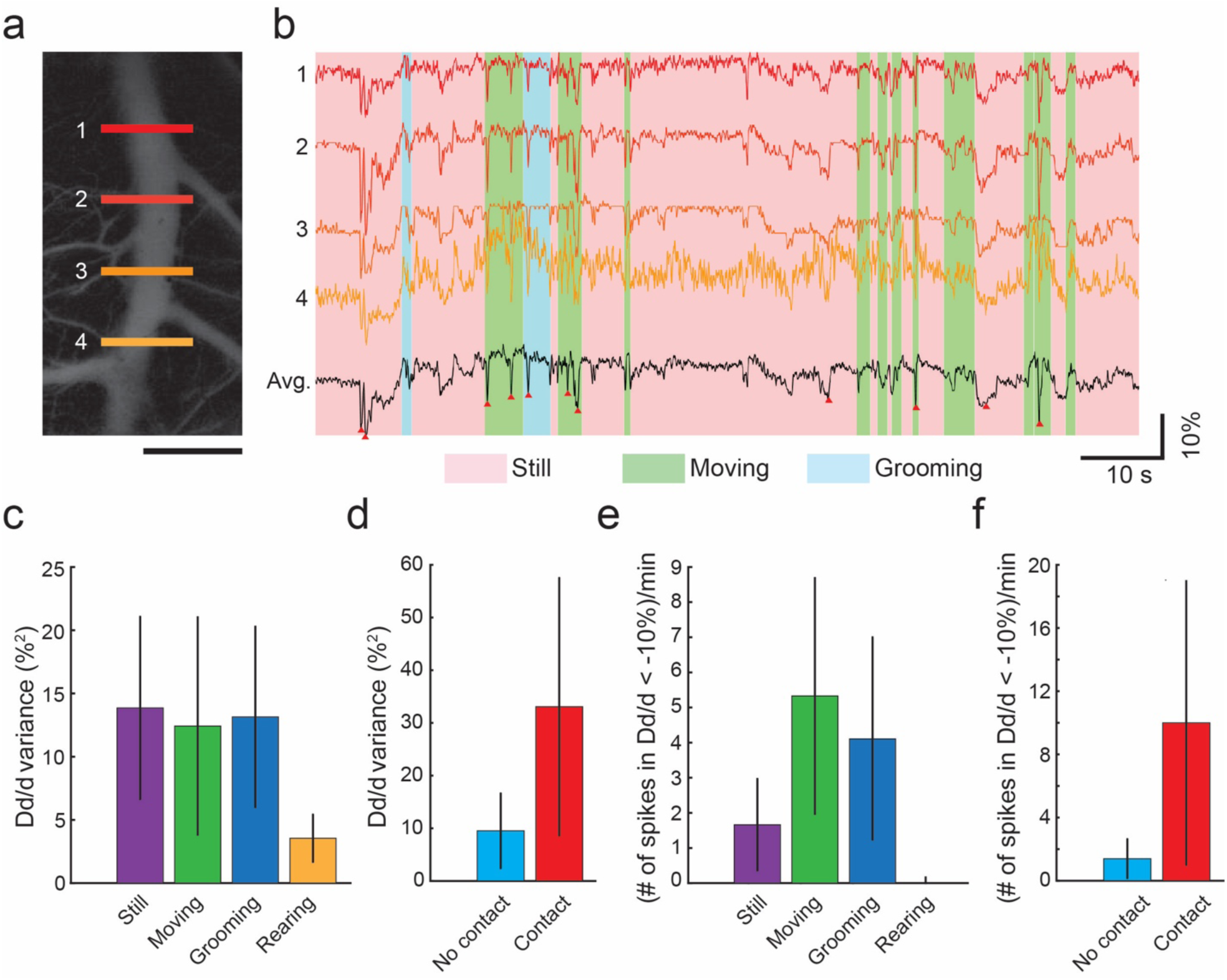
Tracking diameter of superior sagittal sinus (SSS) during free and social behavior: **(a)** Image of SSS in the dorsal cortex. Numbered lines indicate cross sections selected for diameter tracking analysis in **b**. Scale bar, 1 mm. **(b)** Time series evolution of percentage change of vessel diameter at the cross sections specified in **a** and their average during still, moving, and grooming behaviors. Red triangles along the average trace indicate spikes exceeding 10% prominence. **(c)** Bar graph of the variance in percentage change of vessel diameter (n = 6 trials, n = 3 mice) by behavior. **(d)** Bar graph of the variance in percentage change of vessel diameter by social activity (n = 3 trials, n = 3 mice). **(e)** Bar graph of the rate of occurrence of 10% prominence spikes by behavior (n = 6 trials, n = 3 mice). **(f)** Bar graph of the rate of occurrence of 10% prominence spikes by social activity (n = 3 trials, n = 3 mice).

### Mapping cortical functional connectivity during open field exploration and social interactions

Mesoscale imaging has been used to reveal functional connectivity between cortical areas during spontaneous resting state activity^6,16^. Such functional connectivity studies have thus far been conducted in head-fixed animals^6,21^. Here we used the Mesoscope to examine functional connectivity during open field behavior of solitary mice and during social interactions with a companion mouse. An overhead camera tracked behavior simultaneously during Mesoscope imaging (**Fig. 6a**). This was followed by analysis of the behavior videos using DeepLabCut^42^ (**Fig. 6b)**. Solitary open field trials, each lasting 2 minutes, were manually segregated into four types of behaviors – periods when mice were still, moving, grooming, or rearing. On average, mice spent 70.6% ± 9.0% of the time remaining still (n = 6 trials, 3 mice), while spending 20.7% ± 7.51% of time moving within the cage. Grooming and rearing were much less frequent and shorter in durations, accounting for 7.7% ± 6.02% and 1.04% ± 1.17% respectively (**Fig. 6c**). To study social behavior, mice bearing the Mesoscope were allowed to first explore the arena for 4 minutes before a companion mouse of the same sex was introduced (n = 3 trials, 3 mice). Social trials lasted a further 2 minutes. Mice spent 41.305% ± 19.5% of time socially interacting with each other, including touching whiskers or the body (**Fig. 6d**).

**Figure 6:**
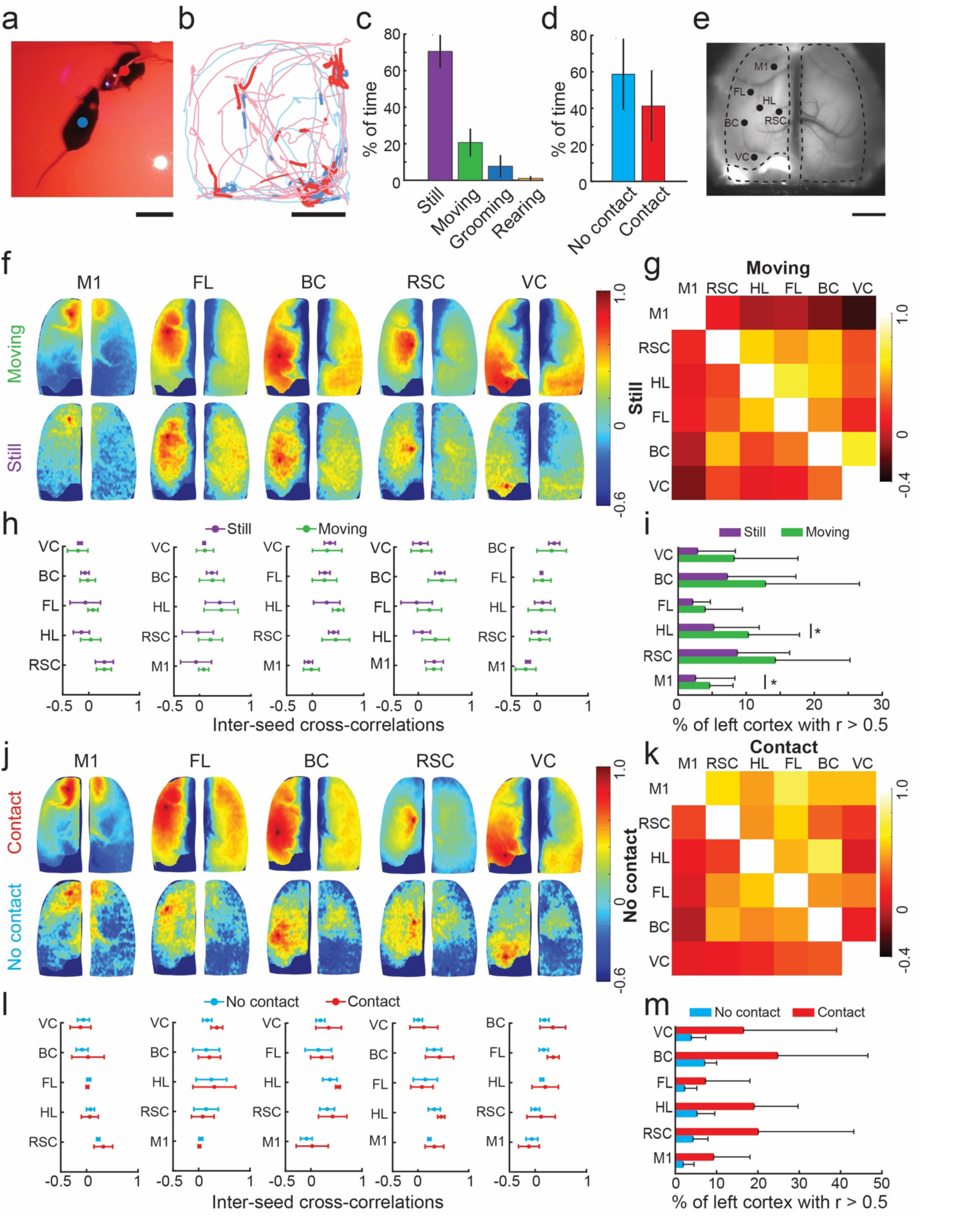
Mesoscale imaging of cortex during free and social behavior: **(a)** Photo of a male GCaMP6f mouse fitted with a Mesoscope interacting with a male C57BL6 mouse in a 33 cm square open field arena. Mesoscope attached to mice annotated with red marker. Companion mouse annotated with blue marker. Scale bar, 5 cm. **(b)** Trace of instantaneous locations of the two mice shown during a 4-minute trial. The blue trace indicates a mouse with a Mesoscope fitted, and the red trace indicates the companion mouse. The segments of darker blue and red indicate positions of the mice when they were within 5 cm of each other. Scale bar, 10 cm. **(c)** Bar graph showing the average percentage of time spent still, moving, grooming, and rearing during open field behavior (n = 6 trials). **(d)** Bar graph showing the average percentage of time spent in contact and not in contact with the companion mouse behavior during trials (n = 3 trials). **(e)** Representative image of the cortex captured by the Mesoscope during a social behavior trial. Locations of the seed pixels used for analysis in **f-h** and **j-l** are indicated. Scale bar 2 mm **(f)** Seed pixel correlation maps of selected ROIs: motor cortex (M1), frontal lobe (FL), barrel cortex (BC), retrosplenial cortex (RSC), visual cortex (VC) and hind limb (HL) during movement and no movement. *Top row:* Average of cross correlations during 1 s epochs of movement. *Bottom row:* Average of cross correlations during 1 s epochs of no movement. **(g)** Matrix of changes in correlation between selected ROIs in **f** during still and moving behavior epochs. The bottom triangular section represents no motion. The top triangular section represents motion. **(h)** Line plots of average inter-seed cross correlations (n = 11 trials, n = 3 mice). **(i)** Summary plot of change in percentage of cortical hemisphere with r > 0.5 with respect to seed pixels analyzed. * indicates p < 0.05 (Mann-Whitney U-test, n = 11 trials, n = 3 mice). **(j)** Seed pixel correlation maps of selected ROIs M1, FL, BC, RSC, VC and HL during non-social and social behavior epochs. **(k)** Matrix of changes in correlation between selected ROIs in **i** during no contact and contact with the companion mouse epochs. The bottom triangular section represents no contact. The top triangular section represents contact. **(l)** Line plots of average inter-seed cross correlations (n= 3 trials, n = 3 mice). **(m)** Summary plot of change in percentage of cortical hemisphere with r > 0.5 with respect to seed pixels analyzed.

We next constructed seed-pixel cross-correlation maps of the cortex from the calcium activity videos acquired by the Mesoscope during open field behavior. Maps with respect to six seeds within the motor cortex (M1), the forelimb (FL), the hind limb (HL), the barrel cortex (BC) areas in the somatosensory cortex, the retrosplenial cortex (RSC) and the visual cortex (VC) were analyzed in each mouse (**Fig. 6e**). Representative seed pixel correlation maps with respect to 5 seeds from a mouse during an open field behavior trial are illustrated in **Figure 6f.** Correlations between the seed locations changed when the animal was moving versus still. Between the seeds, correlations increased, particularly between those located within somatosensory cortex (**Fig. 6g**). Overall movement induced increased variance in the inter-seed correlations (**Fig. 6h**). The area of the left hemisphere of the cortex that is highly correlated with respect to a given seed location, increased for all seeds analyzed when the animal was moving (**Fig. 6i**) and these increases were found to be significantly higher for M1 and HL (n = 11 trials, 3 mice, p < 0.05, Mann-Whitney U-test). Similarly, seed-pixel correlation maps were constructed for mice engaging in social behaviors (**Fig. 6j**). Most social interactions involved mice pausing movement to engage in social behaviors (**Fig. 6b**). Despite the lack of movement, intra-cortical connectivity was increased during times where the mice were engaged in social behaviors (**Fig. 6j-m**). These results demonstrate the utility of the Mesoscope to study functional connectivity during two behaviors that are unique to freely behaving mice.

### Imaging glutamate release dynamics during wakefulness and natural sleep

As a final demonstration, we used the Mesoscope to measure dynamic changes in extracellular glutamate release in the cortex during transition from wakefulness to natural sleep. Glutamate is an important neurotransmitter that regulates neural activity and cerebral metabolism during wakefulness and sleep. However, much of the previous work studying extracellular release has been done using fixed-potential amperometry^43^, or optical imaging in head-fixed mice^44^. Inducing sleep in head-fixed mice is challenging and typically requires sleep deprivation which can alter the overall sleep structure and patterns of rapid eye-movement (REM) and non-REM (NREM) sleep^45,46^. The design flexibility of the See-Shells allowed us to incorporate local field potential (LFP) recording electrodes in the dorsal hippocampus in Emx-CaMKII-Ai85 mice, expressing iGluSnFR in glutamatergic neocortical neurons^33,44,47^. Mice were allowed to naturally transition to sleep in their home cage. Montages of glutamate activity across the whole dorsal cortex during wakefulness, REM sleep and NREM sleep are shown in **Figure 7a**. Hippocampal LFP clearly indicated transition to NREM sleep and subsequently REM sleep (**Fig. 7b, c** and **h**). Consistent with previous studies, spontaneous cortical activity patterns during quiet wakefulness and NREM were highly synchronized across hemispheres^48^. In addition, cortical activity patterns were not necessarily global changes in activity state and were instead composed of complex local activity patterns (**Fig. 7a**). The transition from wakefulness to NREM sleep and REM sleep resulted in decreased fluorescence, indicating reduced cortical glutamate activity (**Fig. 7a-d**). We also observed a reduction of slow cortical glutamate fluctuations during REM sleep (**Fig. 7e**). Correlation analysis of cortical activity revealed that connectivity decreases in REM sleep compared to quiet wakefulness and NREM sleep (**Fig. 7f** and **g**). Moreover, consistent with previous studies, the strength of functional connectivity is less in NREM sleep compared to quiet wakefulness^48^. The Mesoscope attachment also allowed us to study the transition from REM sleep to wakefulness, wherein increased glutamate activity across the cortex was observed (**Fig. 7h** and **i**).

**Figure 7:**
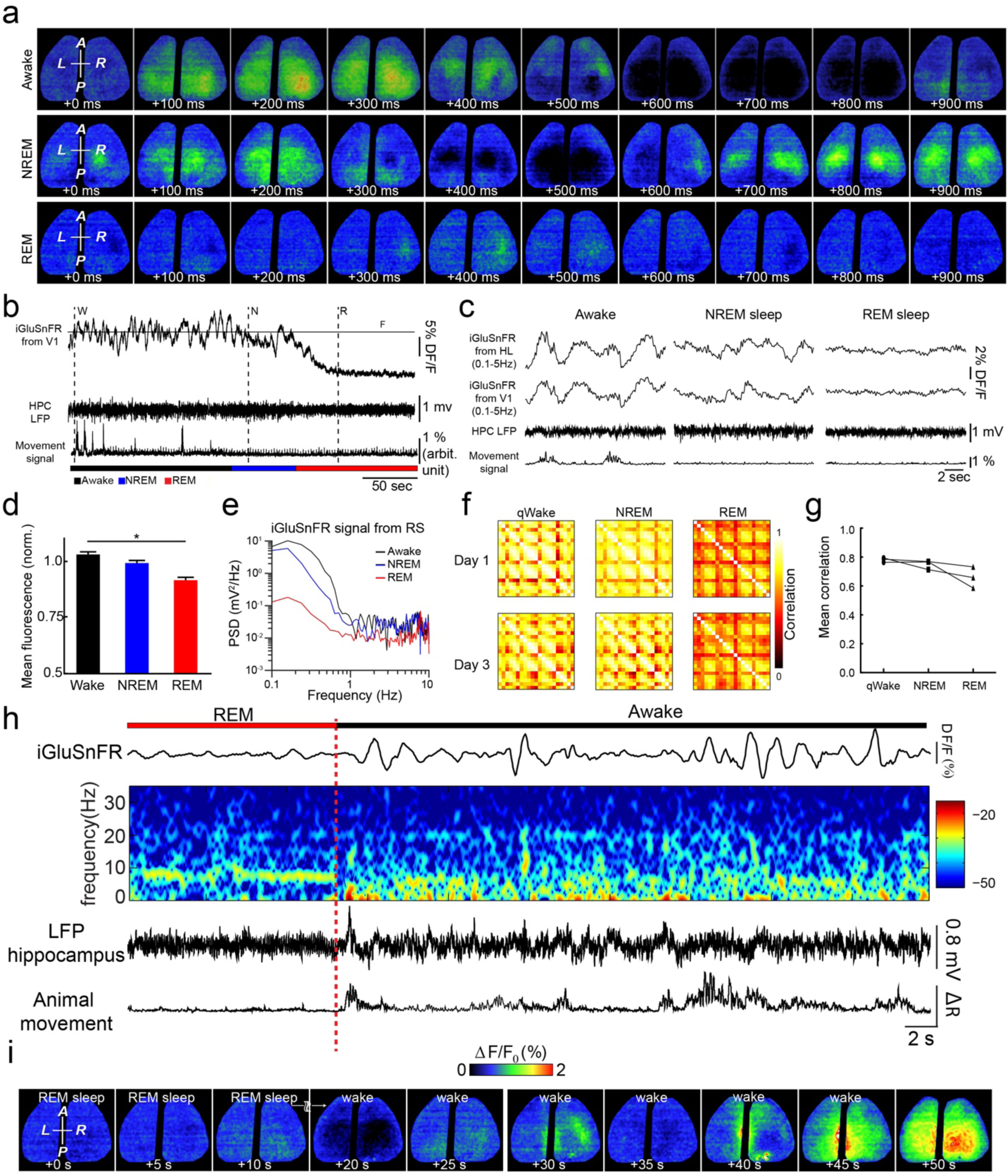
Combined electrophysiological recording and mesoscale imaging of brain activity during wakefulness and sleep: **(a)** Montages of change in glutamate activity over 1s period during awake, REM and NREM sleep states. **(b)** Representative traces of raw glutamate signals from V1, hippocampal LFP, and video-based movement signals during states of waking, NREM sleep and REM sleep in a freely moving mouse. F_0_ is the baseline glutamate signal calculated by averaging the fluorescence over the entire recording time. **(c)** Magnification of glutamate, LFP and movement signals selected in **b. (d)** Grouped mean raw glutamate signal from the entire cortex (normalized to baseline) during wakefulness, NREM sleep and REM sleep (n = 3 mice, Friedman paired nonparametric test; post-hoc multiple comparison with Dunn’s correction * indicates p < 0.05. **(e)** Spectral analysis of glutamate signal shows differences in power content between three states. **(f)** Change in the correlation maps of cortical activity between quiet wakefulness, NREM sleep and REM sleep. Cortex was divided into 21 ROIs. Similar changes between quiet wakefulness, NREM sleep, and REM sleep can be observed on different days. **(g)** Reduction of mean correlation of cortical activity from waking and NREM sleep to REM sleep (n = 3 mice). **(h)** Change in glutamate activity, hippocampal LFP and animal movement during transition from REM to wakefulness **(i)** Montage of cortical glutamate changes during the transition shown in **h**.

## DISCUSSION

We have introduced a new neurotechnology for mesoscale activity mapping of the dorsal cortex in freely behaving mice. The Mesoscope represents a significant addition to toolkits available for neuroscientists to study how multiple regions of the cortex coordinate their activity to produce behavior. The overall weight of the Mesoscope was comparable to existing head-mounted imaging devices^29^, and it did not significantly affect locomotion in an open field arena. Despite using relatively simple optics and an inexpensive imaging sensor, the Mesoscope performs comparably to a conventional epifluorescence macroscope when used for imaging calcium dynamics across the dorsal cortex. The Mesoscope can simultaneously perform both fluorescence imaging and reflectance imaging to correct for hemodynamic changes. Perhaps the biggest advantage of the Mesoscope when compared to existing preparations for studying mesoscale dynamics is the ability to perform experiments in animals performing complex behaviors that would be impossible to study in head-fixed animals. As demonstrated in this study, we constructed to our knowledge the first functional connectivity maps in mice locomoting and socially interacting with a companion mouse in an open field arena.

Among mammalian models used in neuroscience, mice have the widest range of transgenic animals for broad expression of genetically encoded calcium indicators^34,49,50^, voltage indicators^5^ as well as reporters of neurotransmitters^4^. In this study we also take advantage of available transgenic mice expressing reporters of glutamate to study dynamics glutamate release in the cortex in naturally sleeping mice. Combined with a wide range of mouse models of neurodegenerative and neuropsychiatric disorders^51^, the Mesoscope should enable studies of mesoscale cortical activity mediating a wide range of complex behaviors in healthy mice and how these activities may be disrupted in diseased states.

The Mesoscope uses simple hardware to enable simultaneous imaging of a large area of the dorsal cortex. Once the custom fabricated parts are procured, the Mesoscope can be rapidly assembled (< 1 day). The design takes advantage of the sensors made available through the open source miniature imaging systems pipeline^27^. This is a rapidly evolving ecosystem where new sensors with increased sensitivity and imaging speed^52^, miniaturization^53^, wireless imaging capabilities^54^ being developed. Relatively simple modifications to the current Mesoscope design will allow these newer sensors to be incorporated.

The Mesoscope performs reflectance measurements at a single wavelength using a green LED to correct for hemodynamic effects. Multiple wavelength reflectance measurements allow for a more accurate correction of hemodynamic effects^38,55^. In future versions, an additional red LED could be incorporated to obtain reflectance measurements at two wavelengths. This will allow more accurate corrections for hemodynamic effects. Alternately, issues with hemodynamic corrections could be addressed by including illumination at 405 nm to excite GCaMP6f at its isosbestic point^56^.

The Mesoscope architecture can also be readily adapted to image red-shifted fluorescent reporters where hemodynamic effects are not prevalent^2,57^.

The Mesoscope was designed to be compatible with our previously developed See-Shells^41^. Compared to intact skull preparations, implanting the See-Shells increases the surgical complexity. However, the Mesoscope images the cortex at higher resolution compared to intact skull imaging. This allows us to readily perceive any motion that may cause artifacts in the imaging. For instance, we tracked minute changes in SSS diameter and were able to precisely eliminate locations close to the SSS where such motion artifacts were observed from further analysis. Higher resolution images can be used to identify surface blood vessels where lateral motion artifacts and hemodynamics cannot be corrected for, and the surface blood vessels can be digitally masked prior to analysis. Further, the digital design methodology of the See-Shell could be used to develop polymer skulls with an expanded FOV to encompass the cerebellar cortex or more lateral regions of the cortex. Finally, future Mesoscopes could be designed to incorporate miniaturized amplifiers for integrating chronically implanted recording electrodes for simultaneous mesoscale imaging and deep brain neural recordings, or to interface with electrodes incorporated in the See-Shells for electrocorticography (ECoG)^58^.

## METHODS

### Mesoscope design fabrication and assembly

#### Design

The Mesoscope was designed using a computer-aided design (CAD) program (SolidWorks, Dassault Inc.). The top and bottom housing were CNC milled from Delrin. Smaller components, including the filter cubes, were 3D printed using a desktop stereolithography (SLA) printer (Form 2, Formlabs Inc.) with UV-curable black polymethylmethacrylate (PMMA) resin (RS-F2-GPBK-04, Formlabs Inc.). The excitation light sources (blue and green channels) and collector lens were mounted in the bottom housing of the Mesoscope. A custom diced bandpass excitation filter (450-490 nm, 3 mm × 3 mm × 1 mm, ET 470/40x, Chroma) was bonded to the bottom face of the cube using UV-curable optical glue (AA 352 Light Cure Adhesive, LOCTITE). A blue LED (LED LUXEON Rebel Color Blue 470 nm, Digikey Inc.) was bonded to the top face of the filter cube. Three such filter cubes were assembled, and the three blue LEDs were serially wired using 29-gauge wires (Low-Voltage High-Temperature Black, Red Wire with FEP Insulation 29 Wire Gauge, McMaster). Additionally, 1.5 mm x 2 mm x 1 mm copper plates were soldered to the large pad on the backs of the LEDs to act as heatsinks. UV-curable black resin was applied to the back of each blue LED to encapsulate the wires for stabilization. The three excitation cubes were then inserted into the excitation light source slots in the bottom housing. A green LED (LED Lighting Color LUXEON Rebel Color Green 530nm, Digikey Inc.) was bonded to the green LED slot on the bottom housing using cyanoacrylate glue (Professional Super Glue, LOCTITE). Two circular neodymium magnets (4 mm diameter x 1 mm, B07C8ZZ2K9, Amazon) were bonded to the bottom surface of the bottom housing with cyanoacrylate glue. A biconvex collector lens (3 mm diameter x 4.5 mm Achromatic Doublet Lens, Edmund Optics) was gently inserted and press fit into the circular slot at the top of the bottom housing, followed by mounting a custom diced bandpass emission filter (500 - 550 nm, 4 mm × 4 mm × 1 mm, ET525/50m, Chroma). This completed the assembly of the bottom of the Mesoscope. A CMOS sensor (Miniscope CMOS PCB, Labmaker) was fastened to top housing using M1 Thread-Forming Screws (96817A704, McMaster Carr). The top housing was slid onto the rectangular shaft of the bottom housing and the focusing was adjusted and then fixed using 316 Stainless Steel 0-80 screws (91735A262, McMaster Carr).

#### Wiring

A single coax cable was used to connect the CMOS sensor to the main DAQ board. Images were acquired at 30 frames per second (fps). For synchronized illumination of alternate frames with blue and green light, the external trigger output from the CMOS sensor DAQ was sent to a microcontroller (Teensy 3.5, PJRC). At each odd frame, a microcontroller sent a 3.3V transistor-transistor logic (TTL) pulse to a power metal oxide field effect transistor (MosFET IRL 520, Digikey) relay to turn on the three blue LEDs for 20 ms. At each even frame, a second TTL pulse lasting 4 ms was sent to a dedicated MosFET powering the green LED. The wires powering the LEDs were routed through a circular hole in the top housing and then a commutator (Carousel Commutator 1x DHST 2x LED, Plexon) for strain relief. High wattage resistors were used in the switching circuit (**Supplementary Fig. 2**) and the power supplies for each illumination source were used to modulate the current delivered to the LEDs and tune light illumination output during *in vivo* experiments.

#### See-Shell and protective cap preparation

The See-Shell implant modified for quick fixing the Mesoscope was assembled using the technique adapted from our previous work^35^. Briefly, the frame of the See-Shell was 3D printed using a desktop SLA printer with UV-curable black PMMA resin. A 50 µm thick PET film (Melinex, Dupont Inc.) was bonded to the PMMA frame using quick setting epoxy (ScotchWeld dp100 Plus Clear, 3M Inc.). Two circular neodymium magnets were inserted into the slots of the lateral tabs and fixed using cyanoacrylate glue. An 0-80 nut was inserted into the hole in the posterior tab of the implant (Brass Hex Nut, Narrow, 0-80 Thread Size, McMaster). The hole in the posterior tab was briefly heated using a solder iron and an 0-80 nut was press fit into the hole. Similarly, the protective cap was 3D printed using an SLA printer with black PMMA resin, and two circular neodymium magnets were mounted into the slots of the bottom surface of the cap.

### Surgical Implantation

#### See-Shell implantation

All animal experiments were conducted in accordance with protocol approved by the University of Minnesota’s Institutional Animal Care and Use Committee (IACUC) and the University of Lethbridge’s Animal Care Committee. Transgenic mice expressing sensors for calcium (Thy1-GCamp6f mice^34^) or glutamate (Emx-CaMKII-Ai85^44^) were administered 2 mg/kg of slow release Buprenorphine (Buprenorphine SR-LAB, Zoopharm Inc.) and 2 mg/kg of Meloxicam for analgesia and inflammation prevention, respectively. Mice were anesthetized in an induction chamber containing 3-5% isoflurane in pure oxygen. Once the mice were anesthetized, isoflurane was reduced to 1-2% and hair was removed from the scalp using a trimmer and saline-soaked cotton tipped applicators. Sterile eye ointment (Puralube, Dechra Veterinary Products) was applied to the eyes to prevent drying. Mice were then transferred and affixed to either a standard rodent stereotax (Model 900LS, Kopf Inc.) or an automated robotic surgery platform, the Craniobot^59,60^, both of which had heating pads to maintain body temperature The scalp was sterilized by alternately scrubbing with 10% w/v Povidone-Iodine solution (Betadine, Avrio Health L.P.) and 70% ethanol three times. The scalp above the dorsal cortex was excised using surgical scissors and fascia was removed from the skull surface by repeatedly rubbing the skull with a 0.5 mm micro-curette (10080-05, Fine Science Tools). A large craniotomy over the dorsal cortex was performed either manually using a high-speed dental drill or automatically with the Craniobot. The craniotomy was performed taking great care to ensure the underlying dura remained intact and no damage was caused to the brain. Once the skull was removed, the exposed brain was immediately covered in a gauze pad soaked in sterile saline to keep the brain hydrated.

The assembled See-Shell implant was sterilized by briefly (1-2 minutes) soaking in 70% ethanol, followed by thorough rinsing with sterile saline. The gauze pad on the brain was removed and excess blood from the craniotomy was cleared using pointed cotton tip applicators (823-WC, Puritan). The See-Shell implant was gently placed on the skull, and the area of the skull surrounding the See-Shell was dried using cotton tipped applicators. A few drops of surgical glue (Vetbond, 3M Inc.) was applied around the edge of the See-Shell followed by opaque dental cement (Metabond, Parkell Inc.) to adhere it to the skull. The cement fully cured before the protective cap was magnetically attached. Mice recovered on a heated recovery pad (72-0492, Harvard Apparatus Inc.) until fully ambulatory before returning to a clean home cage. Post-operative analgesia and anti-inflammatory drugs were administered for up to 72 hours post-surgery. Mice recovered from surgery for at least 7 days before experimentation.

#### Electrophysiology

In experiments where we performed simultaneous electrophysiology and mesoscale imaging, the hippocampal electrode was implanted through a hole outside the area of the See-Shell craniotomy. Briefly, a bipolar (tip separation of 500 µm) electrode made from Teflon-coated stainless-steel wire (bare diameter 50.8 µm) was placed in the pyramidal layer of the right dorsal hippocampus. It was inserted posterior to the occipital suture at a 33-degree angle (with respect to the vertical axis) according to the following coordinates relative to bregma: ML: 2.3 mm; dorsoventral (DV): 2.2 to 2.5 mm. The position of the electrode tips was confirmed using an audio monitor (AM8C #72×32B, Grass Instrument Co.). After electrode implantation, the craniotomy was performed and the See-Shell was implanted on the skull as described above.

#### Reinforced, intact skull preparation

Intact skull surgical preparation followed the protocol described previously^36^. The animal was aseptically prepared, followed by removal of the scalp and fascia from the skull as described above. A skull anchor screw was implanted over the occipital bone. Clear dental cement (S399, Parkell Inc.) was applied over the surface of the skull and a sterilized 8 mm diameter circular glass coverslip (Deckgläser, Marienfeld-Superior Inc.) was pressed on top of the dental cement. Care was taken to ensure no air bubbles were present in the FOV. A small titanium headpost was then attached using radiopaque dental cement.

### Benchtop characterization optical imaging performance of the Mesoscope

#### Resolution testing

To measure the resolution, a transparent 1951 USAF test target was cut to isolate each group and placed onto a See-Shell window. Seven volts was provided to the three blue LEDs to illuminate the target. The center of the target was placed onto a 3D printed inverse mold of the See-Shell contour with markings for 8 different test locations in the FOV. A See-Shell implant was pressed onto the inverse mold with the target and images were taken in each location (**Fig. 1e**).

#### Illumination profile

To measure the uniformity of illumination, 3% agar gel consisting of 10% concentration fluorescein dye (F2456, Sigma Aldrich Inc.) was prepared. A custom 3D printed acrylic container was filled with fluorescein dye infused agar gel phantom. The See-Shell implant was placed on the container such that the entire bottom surface of the See-Shell was uniformly coated with fluorescent gel phantom. The Mesoscope was attached to the See-Shell using the magnetic interlocking mechanism. Single images were acquired using the Mesoscope, and the current delivered to the LEDs was modulated using power supplies in the switching circuit (**Supplementary Fig. 2**) to eliminate FOV saturation. The captured images were analyzed in MATLAB (MathWorks, Inc) using custom code.

#### LED switching dynamics and LED power stability testing

To test the stability of the blue LEDs across multiple cycles, they were turned on for 10 ms and light output was measured using a photoresistor (NSL-19M51, Advanced Photonix). For a total of 6 trials, 130 cycles were collected per trial and the voltage output of the photoresistor was analyzed in MATLAB. To ensure light bleeding was minimized between blue and green frames and to determine a pulse duration for each LED type, experiments were performed on fluorescein dye infused agar phantom where blue LEDs were pulsed while green LEDs remained off. For each frame, the mean intensity across the FOV was quantified (**Supplementary Note 2**).

### Focusing and calibration of Mesoscope

Before every experiment, mice were lightly anesthetized (0.5-1% isoflurane in pure oxygen) and head-fixed in a stereotax to remove the protective cap and clean the See-Shell surface of any debris using distilled water and cotton tipped applicators. The Mesoscope was placed on the implant and secured via the magnetic interlocking mechanism and an 0-80 screw in the posterior tab. The green LED was switched on and the position of the top housing relative to the bottom housing was manually adjusted until the area around the midline was in focus and the adjustment screw was tightened to secure it in place. Once focused, the three blue LEDs and green LED were alternately pulsed, and their intensities were adjusted by modulating the power delivered to the LEDS using each power supply. (**Supplementary Fig. 2**).

### *In vivo* calcium imaging in anesthetized mice

Initial *in vivo* calcium imaging experiments were performed in anesthetized animals to compare imaging capabilities to a conventional macroscope and imaging sensory evoked responses. Mice were lightly anesthetized and affixed to a custom stereotaxic setup under a conventional epifluorescence macroscope (Leica MZ10F, Leica AG). Images from the macroscope were captured using a sCMOS camera (Orca Flash 4.0, Hamamatsu Inc.). Bright field images were first captured to assess the quality of the cranial window followed by epifluorescence imaging. Since blue illumination was provided every alternate frame in the Mesoscope, the effective image capture rate was 15 Hz. To closely replicate these imaging conditions, the sCMOS camera was configured to acquire images at 8 bits and 30 fps. In the final imaging analysis, every other frame was discarded to ensure an effective acquisition rate of 15 Hz. The Mesoscope was attached immediately after image acquisition from the macroscope and 4 minutes of video was captured at 30 fps. To measure sensory evoked responses, mice were lightly anesthetized. To evaluate the response to vibrational stimuli, a 1 s vibrational stimulus was provided to the hindlimb using a 3V DC mini vibration motor (A00000464, BestTong) at 100 Hz. For visual stimulus, a white LED was positioned ∼ 2 cm from the left eye of the mouse to cover the fovea in the visual field. Hundred ms flashes of light were presented after ensuring light from the LED was shielded to confine stimulus delivery to the left eye.

### Open field and social behavior experiments

Mice recovered from surgery for at least 7 days before experimentation. The recovery period was followed by an acclimatizing period of 3-5 days in which a single experimenter handled each mouse for 5-15 minutes. Mice were then introduced into a 33 cm square open field behavioral arena and allowed to freely locomote, while behavior was recorded using an overhead camera (ELP-USB8MP02G-L75, ELP). To evaluate the effect of the Mesoscope on open field locomotion, five mice in each group (non-surgical control mice, mice implanted with See-Shells, and mice implanted with See-Shells bearing a head-mounted Mesoscope) performed trials lasting five minutes. Trials were repeated for each mouse over five consecutive days, totaling 25 trials in each group. For open field and social behavior imaging experiments, the mice were fitted with a Mesoscope replica with the same weight as a fully assembled Mesoscope during the initial handling period. Before every experiment, mice were lightly anesthetized and head-fixed in a stereotax to remove the protective cap and clean the See-Shell surface of any debris using distilled water and cotton tipped applicators. The Mesoscope was fitted onto the mouse and the top housing was adjusted to focus the area around the midline. The mice were quickly transferred to an open field arena and allowed to regain full consciousness for at least 10 minutes following complete transition to wakefulness. Typically, the experimental setup required waiting much longer than 10 minutes after anesthesia was withdrawn. Experiments were conducted for 6 minutes each, including a two-minute period to allow the LEDs to warm up to their maximum intensity. During social behavior experiment trials, a C57BL/6 mouse of the same sex was gently introduced into the arena by an experimenter 4 minutes after initiation of the trial. After the LED warm up period, this allowed 2 minutes of imaging of the solitary mouse and 2 minutes with a companion mouse.

### Sleep recording experiment

Mice recovered from surgery for at least 7 days and then habituated for 7 days in the recording setup with the Mesoscope mounted on their head. Each mouse was recorded for two sessions each lasting two hours. Hippocampal LFP was amplified (×1000) and filtered (0.1-300 Hz) using a Grass P5 Series AC amplifier (Grass Instrument Co.) and was sampled at 1 kHz using a data acquisition system (Axon Instruments). A camera (Camera Module V2 #E305654, Raspberry Pi) was used to record behavior during the recording.

### Data Analysis

#### Behavior video analysis

All videos of mouse behavior were captured in .avi format. To quantify open field locomotion of solitary mice, videos were analyzed using Zebtrack^61^ software in MATLAB. Each trial was manually verified to ensure accuracy or correct tracking errors. The total distance travelled was calculated by scaling the distance moved in the arena across the entire 5-minute duration. Mean speed was calculated by dividing the total distance by 5 minutes. The maximum speed was determined by finding the maximum of the norm of the absolute values of the discrete derivatives of the x and y positions of the mouse. A 17 cm square area was defined at the center of the arena as the ‘open field’, and we quantified for how long mice spent time in this area. All statistical comparisons between groups were carried out using Student’s t-test to determine any effects of the Mesoscope on free behavior.

In the social behavior trials, DeepLabCut^42^ was used to track the position of the solitary mouse and the companion mouse over time. Twenty frames from each social behavior trial were randomly extracted using k-means clustering and used as training data for the machine learning algorithm. Four body points were tracked on each mouse via deep neural networks embedded in the DeepLabCut software. The positions of all points were averaged to acquire a center-of-mass-like position of each mouse, which was tracked across all frames to generate the plot shown in **Figure 6b**. To segment the behavior based on different behavior epochs, we manually scored behavior with 1 s precision based upon whether the mouse with the Mesoscope was moving, still, grooming or rearing, and whether the mice engaged in social behavior or not indicated by touching, sniffing, or close proximity to the companion mouse and categorized into “contact” and “no contact” epochs. This data was further processed only if there was consensus among at least 3 of the 4 independent scorers.

#### Imaging data pre-processing

All data from the CMOS sensor was captured in .avi format (RGB, 480×752 image size) and contained alternately pulsing blue and green channel data. The video sequences were processed through a custom MATLAB script. Videos were converted from RGB to grayscale and segmented to exclude the 2-minute LED warmup period at the start of each trial. The resulting video sequence was segmented into individual blue and green channel videos.

#### Sensory evoked responses

The blue and green channel videos contained 18 stimulus trials per experiment. A time stamp file generated during the experiment was used to segment this video into its respective 18 trials. We detrended and spatially filtered each individual trial using a custom weighted spatial filter to remove additional illumination artifacts from the LEDs. The resulting DF/F data for each trial was saved to a .csv file and then averaged into a single .csv DF/F file.

#### Open field experiments

The blue and green channel videos acquired from freely moving animals were additionally corrected for lateral motion artifacts using MoCo rigid motion correction40. A macro was used to automate the process and maintain consistency of MoCo parameters across trials. The first frame of the video was used as the template for motion correction. Parameters used for the MoCo plugin were w = 40 and a down sample value = 0.5. Log files of pixel displacements were saved to .csv files. Resulting videos were saved as .avi files with JPEG compression at 15 fps. The motion corrected blue and green channel videos were re-concatenated back into an alternately pulsing video to mitigate data storage issues during further analysis.

The time stamp data from the Miniscope software contains a list of frames captured by the behavior camera and CMOS sensor. To mitigate frame pacing issues with the CMOS sensor, only frames within ±10% of the specified frame rate were kept. To address the frame dropping issues with the behavior camera (usually dropped ∼400/10800 frames), common frames that exist in the CMOS video and behavior camera video were kept. The resulting data was compared to the scoring data to find the intersect between these frame data sets. Finally, this data set was searched through to find points with paired blue and green frames (for hemodynamic corrections) and served as the final blue and green channel frame data used for analysis.

Each motion corrected pulsing video was binned using a bilinear spatial binning algorithm in MATLAB. Pixels in each channel were separately corrected for global illumination fluctuations using a global correction algorithm^6^ and were spatially filtered using a custom weighted spatial filtering algorithm.

#### Hemodynamic correction

After correction for global and illumination artifacts, the corrected frames were passed to a hemodynamic correction algorithm^13^ shown below in Equation 1:

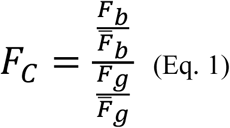

*F*_*e*_ is the corrected fluorescence trace for a blue frame and 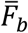 is the mean fluorescence trace across all blue frames in the time series. The green frames’ normalization is conducted similarly. Dividing both the normalized blue and green intensities in every pixel and frame yields *F*_*c*_, the hemodynamic corrected pixel data for every frame in the FOV.

#### Vessel diameter analysis

Vessel diameter tracking was performed using a custom written macro in Fiji. Blue channel videos were inverted, and the background was subtracted using a rolling ball radius of 30 pixels. T rectangular ROI was then drawn around the vessel to be analyzed. The rectangle was used as a guide to create 6 evenly spaced, horizontal lines along the height of the rectangle. The lines were 5 pixels thick and as long as the width of the rectangle. These lines were used as ROIs to track the vessel diameter throughout a trial (**Fig. 5a** and **b**). To calculate vessel diameter, the intensity profile along a line was first plotted. Then, the maximum intensity and its corresponding location in pixels along the line was found. Using half of the maximum intensity as a threshold, the macro then searched for the pixel with the lowest intensity greater than the threshold in both directions from the maximum intensity. The difference between the pixel locations of these two values was then stored as the vessel diameter. This process was repeated for each line and each frame in a video. Values were saved in a .csv file for further analysis.

Post-processing and plotting of data were performed in MATLAB. The percent change in vessel diameter (Dd/d) was calculated for 4 ROIs. The baseline was the average vessel diameter across time at each ROI. Time-series of diameter changes were then plotted as 5-point moving averages along with the average of those traces (**Fig. 5b**). Behavior data was plotted using the consensus among scorers and then padding the behavior epochs on either side by ∼0.5 s (7 frames). Spikes with prominence greater than 10% and longer than 1/3 s were detected using MATLAB. Variances shown in **Figure 5c** and **d** were calculated by first pooling the variances from behavior epochs to obtain a variance for each behavior within a trial. The weighted average of these variances was then taken across trials to obtain a single measure for each behavior.

A Wilcoxon signed-rank test was performed on variances from still, moving, grooming, and rearing behavior (**Fig. 5c**). A paired sample t-test was performed on spike frequencies from still, moving, and grooming behavior (**Fig. 5e**). Statistical tests were not performed for social behavior plots due to the small number of samples (n = 3).

#### DF/F calculations

Intensity values from the selected ROIs from the blue and green illumination videos were computed using FIJI.^62^ Further analysis was performed in MATLAB. Change in fluorescence for both GCaMP and reflectance signals were calculated over a baseline average across the whole time series. Correction for hemodynamics was performed using a ratiometric method as described previously^37^.

#### Seed-pixel correlation maps

Cortical regions of interest were selected within each video trial by removing the background and areas with high vasculature (**Fig. 6e**) and seed pixels within 6 regions (motor cortex (M1), frontal lobe (FL), barrel cortex (BC), retrosplenial cortex (RSC), visual cortex (VC), hind limb (HL)) were selected for further analysis. Seed pixel correlation maps were generated using Pearson’s correlation coefficient (PCC) to compare the correlation between a desired seed pixel corrected GCaMP trace and the other pixels in the FOV. The data from the manual scoring in (**Fig 6c and d**) was used to generate a list of frame numbers where the mouse was either moving or still. These epochs of time are discontinuous between behaviors, so a moving window was applied to each continuous epoch of time. The window length was 1 s (15 frames at 15 fps) with a 0.5 s sliding window. The PCC between the seed pixel trace and every other pixel in the desired regions of the cortex was calculated for each window length portion. The mean of all the PCCs was calculated and stored as the PCC values for each pixel in the ROI relative to the desired seed pixel. This process was repeated for each trial and mouse to generate representative seed pixel maps for one trial as shown in **Figure 6f.** Focusing on the PCC between seed pixels in the same representative trial, we generated a cross-correlational map shown in **Figure 6g**, where the bottom triangular data represents the still behavior PCC values and the top triangular data represents the moving behavior PCC values. The PCC value between the chosen seed pixels were averaged across all trials in spontaneous free behavior to generate the inter-seed cross-correlational plots shown in **Figure 6h**. To measure the change in activity across the left hemisphere where the seed pixels were chosen, the percentage of pixels in this ROI with PCC *r* > 0.5 was averaged across trials (**Fig. 6i**). Similar analysis was performed for social interaction experiments.

#### Glutamate data preprocessing

Raw data of spontaneous glutamate activity was preprocessed based on the following steps: First, the time series of each pixel was filtered using a zero-phase bandpass Chebyshev filter between 0.1 and 5 Hz. Then a baseline signal was calculated by averaging all the frames, and the fluorescence changes were quantified as DF/ F × 100, where F is the filtered signal. To reduce spatial noise, images were filtered by a Gaussian kernel (5 × 5 pixels, sigma = 1).

#### State scoring

Behavioral states were scored visually using hippocampal LFP and movement signal in 10 s epochs. Movement signals were calculated using a previously described algorithm^63^. Active wakefulness (aW) was characterized by theta hippocampal activity and high movements. Quiet wakefulness (qW) was characterized by theta hippocampal activity and minimum movement. NREM sleep (N) was characterized by large irregular activity in hippocampus and no movement. REM sleep (R) was characterized by theta hippocampal activity and no movement.

#### Correlation analysis

A uniform meshgrid with ∼1.2 mm distance between its points was laid on the field of view and signal at each ROI (0.2 mm^2^) was calculated. Quiet wake, NREM sleep and REM sleep were scored for each recording based on above criteria and Pearson correlation coefficients were calculated between each ROIs during qW, NREM and REM. For comparison in **Figure 7f**, correlation matrices were averaged across all ROIs.

## AUTHOR CONTRIBUTIONS

MLR, LG, ML, DS, LG, GJ, SBK designed and engineered the Mesoscope. MLR, DS, JD, ZSN, OH, LG, SBK designed and executed the experiments. MLR, DS, SL, OH, VR and SBK analyzed the data. MLR, DS, SL, VR, JD, ML, SBK wrote the manuscript. MN and MM designed, executed glutamate imaging experiments and analyzed the data and assisted with manuscript writing.

## ACKNOWLEDGEMENTS

SBK acknowledges funds from the Mechanical Engineering department, College of Science and Engineering, MnDRIVE RSAM initiative of the University of Minnesota, Minnesota department of higher education, National Institutes of Health (NIH) 1R21NS103098-01, 1R01NS111028, 1R21NS112886, RF1NS113287 and 1R21NS111196. LG was supported by the University of Minnesota Informatics Institute’s (UMII) graduate fellowship. DS was supported by the University of Minnesota’s Diversity of Views and Experiences (DOVE) fellowship. MLR was supported by 1R21NS103098-01-01S1.

## INSTITUTIONAL APPROVAL

All animal experiments described in this paper were approved by the University of Minnesota’s Institutional Animal Care and Use Committee (IACUC) and the University of Lethbridge’s Animal Care Committee

## DATA AVAILABILITY

Full numerical data shown in the figures are available as a supplementary dataset accompanying the article. Raw imaging datasets will be made available on request.

## CODE AND HARDWARE DESIGN FILES AVAILABILITY

We have made all CAD files for fabricating the Mesoscope available with this article as **Supplementary File 1**. All code associated with operating the Mesoscope will be made available at: www.github.com/bsbrl. We will post updated versions of the software and hardware design files at this location.

## LIST OF SUPPLEMENTARY MATERIAL

1. **Supplementary Figures and Notes**
2. **Supplementary Video 1: Open field behavior of mouse bearing Mesoscope:** Video depicting a mouse bearing a Mesoscope behaving freely in an open field arena. The mouse is free and comfortable to move throughout the entire arena and rear against the arena walls. The mouse appears to be undeterred by the Mesoscope or its associated wiring.
3. **Supplementary Video 2: Stability of mesoscale imaging using the Mesoscope during freely moving behavior:** Video with top view of mouse moving around in the open field. Epochs of 5-10 seconds of different behaviors are shown (locomotion, grooming, and rearing). Inset: Motion corrected video of the corresponding cortical calcium dynamics as captured by the Mesoscope.
4. **Supplementary File 1:** A .zip file archive of computer aided design files used for custom fabricating Mesoscope components.

## Supplementary Figures and Notes

**Supplementary Figure 1:**
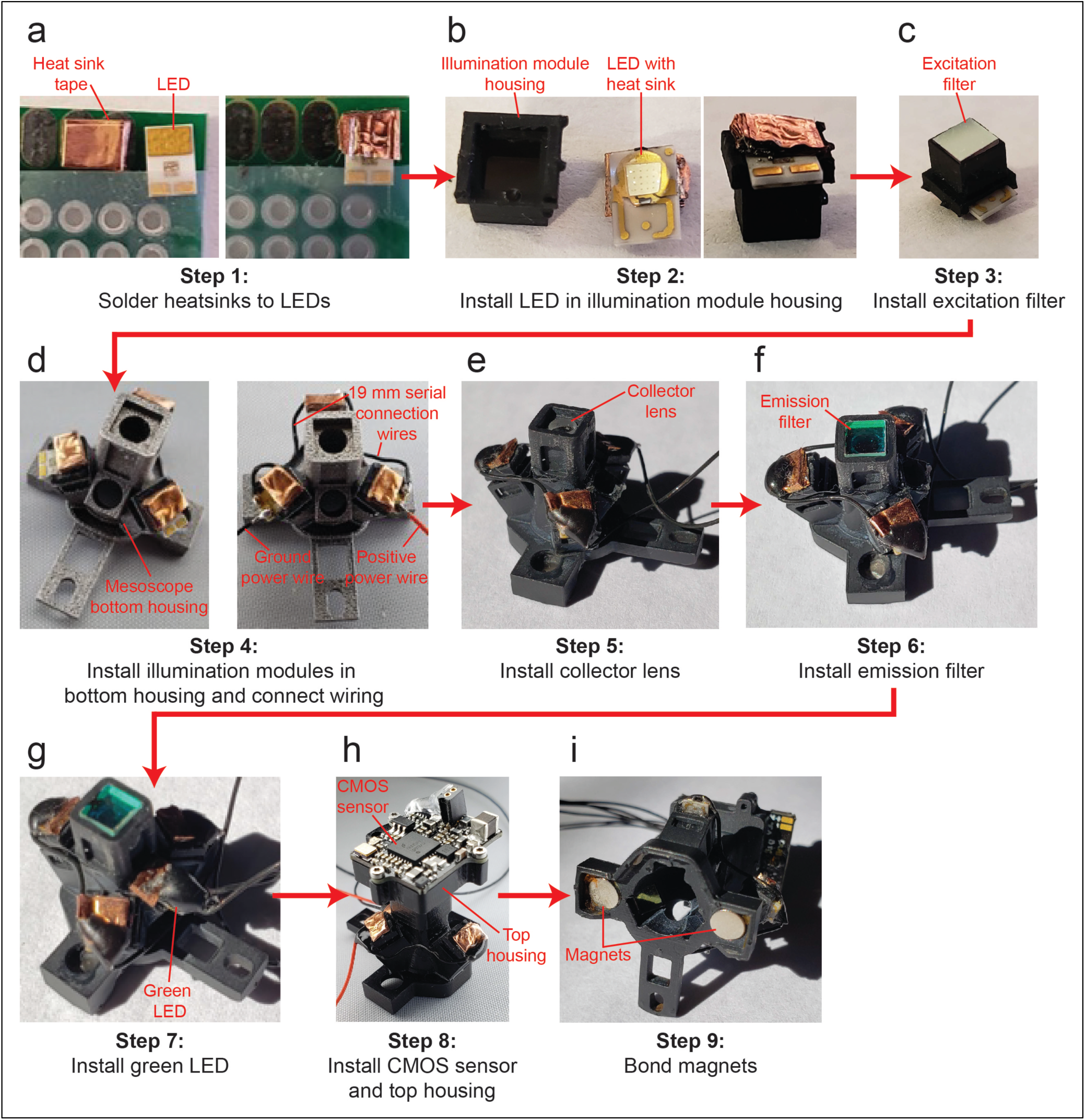
Step by step illustration of Mesoscope assembly.

**Supplementary Figure 2:**
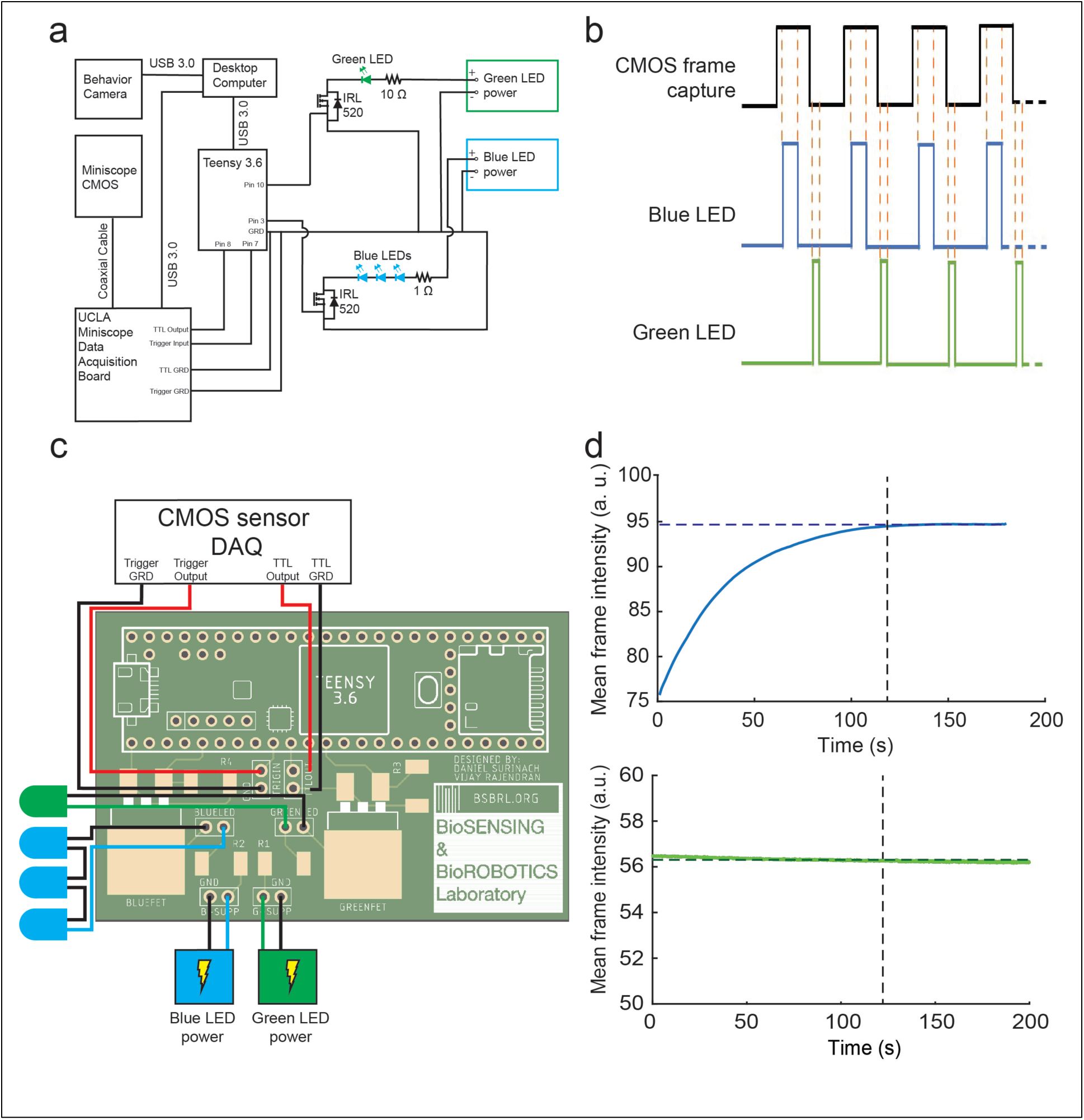
Mesoscope electronics. **(a)** Schematic of circuitry used for alternate illumination of cortex with blue light for fluorescence and green light for reflectance. For every frame captured by the CMOS sensor, the camera data acquisition (DAQ) board sends transistor-transistor-logic (TTL) pulse to a microcontroller (Teensy 3.6). The microcontroller in turn sends alternate TTL pulses to switching circuits that power three blue LEDs and one green LED. The DAQ also controls the behavior camera synchronization with the CMOS sensor to collect cortical and behavioral data simultaneously. **(b)** Frame capture synchronization between the CMOS sensor and LED pulses. At every odd frame, the microcontroller pulses the blue LEDs for 20 ms. At every even frame, the microcontroller pulses the green LED for 4 ms. **(c)** PCB schematic to house the switching circuit. **(d)** Long term LED stability was tested by illuminating a fluorescein dye agar phantom with excitation and emission bands similar to GCaMP6f. The resulting emission signals are collected onto the Mesoscope CMOS sensor. *Top*: The mean frame intensity captured with blue light illumination is shown with the solid blue line. The black dashed line indicates an approximate 120 second warm-up period until a stable illumination point is reached (blue dashed line). *Bottom*: The mean frame intensity data captured with green light illumination is shown with the solid green line. The black dashed line indicates an approximate 120 second warm-up period until a stable illumination point is reached (green dashed line).

**Supplementary Figure 3:**
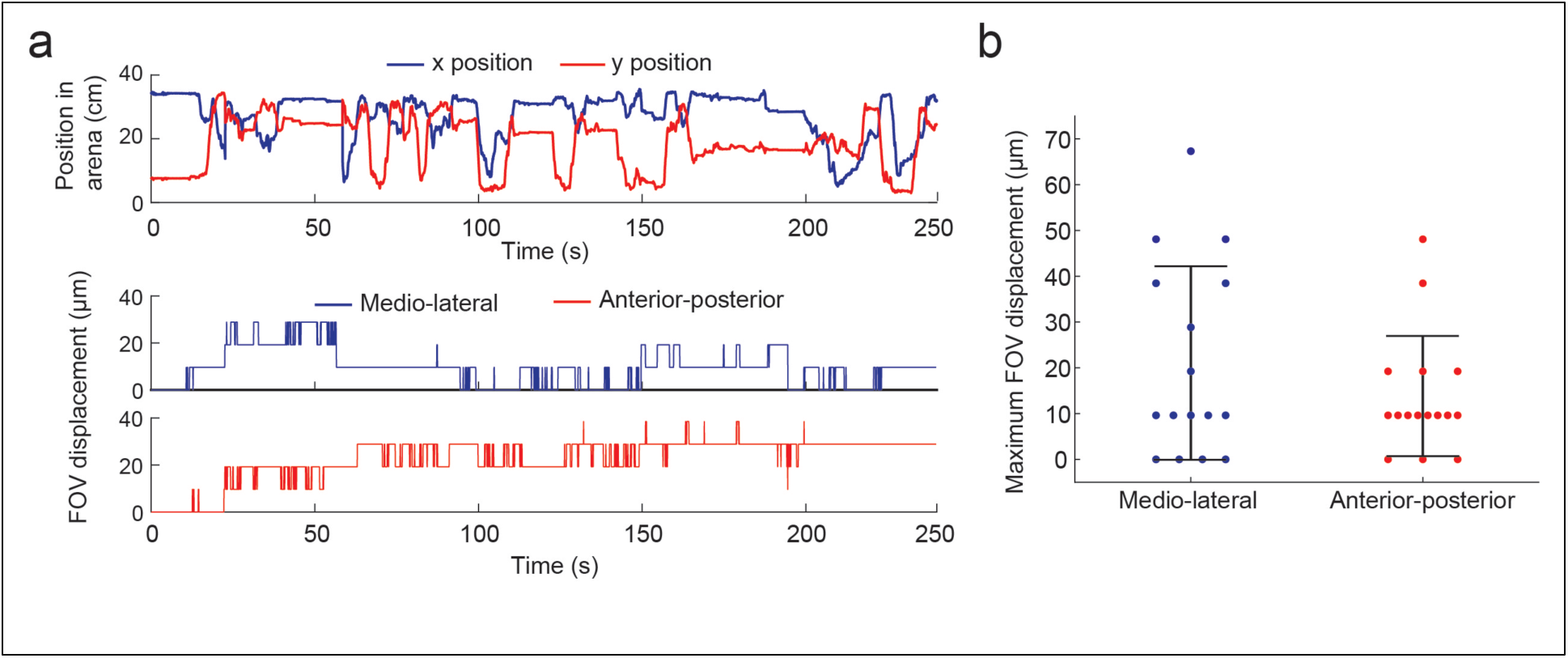
Stability of Mesoscope imaging. (a) *Top:* X and Y coordinates of mouse in a 30cm by 30cm open field arena during a 4-minute trial. *Bottom:* Rigid translations of the Mesoscope FOV to correct for motion artifacts in the calcium dynamics video corresponding to the mouse behaving with the positions depicted in (a). (b) Maximum rigid translations of Mesoscope FOV to correct for motion artifacts in calcium dynamics videos (n = 16 videos).

**Supplementary Note 1: Assembly, wiring and operation of the Mesoscope**

## ASSEMBLY

1. **Pre-fabricate custom parts:** The Mesoscope has custom parts that need to be prefabricated before assembly. These includes the top and bottom housing (**Fig. 1a** and **b**) that are CNC machined out of Delrin, illumination module housings that are 3D printed using a desktop stereolithography (SLA) printer.
1.1. Use filter_cube_ver5.STL (**Supplementary File 1**) to print three illumination module housings. Use files 082019_mesoscope_frame.STL and 10-15-19_frame_cap.STL to print the Mesoscope See-Shells and See-Shells protective cover. Clean the 3D printed parts and post cure under UV light.
1.2. Use files `‘meso4.22_top_delrin.SLDPRT’` and `‘meso4.27_bottom_delrin. SLDPRT’` to CNC machine the top and bottom housing using a 5-axis CNC mill. Alternately, they can be procured from a commercial low volume fabrication services provider (e.g., Protolabs Inc.).
2. **Illumination module assembly:** The illumination module assembly consists of a blue LED, a housing and the excitation filter. Three illumination modules for each excitation light source need to be assembled.
2.1. Solder heat sink to LED: Secure the each LED (1416-1028-1-ND, 1416-1035-1-ND, Digikey) to a prototyping board using tape (**Supplementary Fig. 1a left**).
2.2. Measure out approximately 80mm of heat sink copper tape (B071JKLFXX, Amazon). Mix solder paste and flux in a small container and apply it to the heatsink pad on the LED.
2.3. Fold the heat sink tape 4-5 times so that it is layered, then cut to the dimensions of the heat sink pad (**Supplementary Fig. 1a left)**.
2.4. Press the folded heatsink onto heatsink pad on the LED using forceps. Apply heat to the copper heat sink using a heat gun at 400F, until solder has melted and solidified. Be sure to keep pressure on the copper heat sink so the melting solder can bind to both the LED heat sink pad and the copper heat sink (**Supplementary Fig. 1a right**).
2.5. Repeat this process for each of the three blue LEDs and one green LED.
2.6. Remove the blue LEDs with heatsinks from prototyping board and place on top of the illumination module housing on the top face with the largest square opening and small extrusions to hold the LED in place (**Supplementary Fig. 1b left**). Secure the LED to the filter cube using a few drops of cyanoacrylate glue (1818A5, McMaster) around the perforated edge that the LED fits tightly into. Be sure to avoid getting drops of super glue on the bulb of the LED. Avoid getting drops of super glue on transparent potting covering the diode. Repeat this step for each of the three illumination modules (**Supplementary Fig. 1b right)**.
2.7 Apply a very thin coat of UV-cured optical glue (NOA81, Thorlabs) to the excitation filter (ET470/40x, Chroma) edge and gently insert into the bottom face of the illumination module housing (**Supplementary Fig. 1c**). Use stick tack to handle the filters. Arrows marked on the edge of the diced filters point to the direction of the light propagation. Ensure the filter is inserted in the right orientation. Ensure optical glue does not cover any part of the filter surface. Repeat this step for each of the three illumination modules.
3. **Installing and wiring the illumination module:** Insert the illumination modules into the three illumination arms of the Mesoscope bottom housing (**Supplementary Fig. 1d left**). Cut 2 small pieces of 29-gauge wire to approximately 19 mm (9510T2, McMaster). Cut four 29-gauge wire (2 black and 2 red to provide LEDs power) to the height of the commutator (or power control circuit if no commutator is used). Strip the wires 3-4mm on either side and apply flux to the positive and negative pads of the blue LEDs. Connect the blue LEDs in series using the 19 mm wires. Attach the longer red and black wires to the remaining pads in either terminal of the LEDs wired in series. Check for any shorts in the soldering process. Ensure the LEDs are functioning properly at this step in the assembly. Apply a layer of black epoxy (RS-F2-GPBK-04, FormLabs) to the LED power pads. The epoxy reinforces solder strength at the pad site and provide light shielding (**Supplementary Fig. 1d right**).
4. **Collector lens installation:** Apply optical glue to cylindrical face of the collector lens (47-721, Edmund Optics). Insert into the lens chute within the central shaft of the bottom housing (**Supplementary Fig. 1e**).
5. **Emission filter installation:** Apply optical glue to edges of the emission filter (ET525/50m, Chroma) and insert into the square emission filter slot in the central shaft of the Mesoscope’s bottom housing (**Supplementary Fig. 1f**).
6. **Green LED Installation:** Insert green LED into the remaining circular slot within the bottom housing of the Mesoscope and secure using several drops of super glue. Attach power wires to the positive and negative pads of the LED. Check for any shorts in the soldering process. Ensure the LED is functioning properly at this step in the assembly. Insert green LED into the remaining circular slot within the bottom housing of the Mesoscope and secure using several drops of super glue. Apply a layer of black epoxy to the LED power pads. The epoxy reinforces solder strength at the pad site and provide light shielding (**Supplementary Fig. 1g**).
7. **Top Housing attachment:** Use 18-8 M1 torx flat head set screws (96817A704, McMaster) to fasten the CMOS sensor to the top housing. Slide the top housing on the central shaft of the bottom housing. Use the 0-80 adjustment screw (91735A262, McMaster) to secure the top housing in place. Feed the LED power wires through the wire holder in the top housing. Use a little epoxy to fix the wires in place, ensure some slack to ensure strain relief (**Supplementary Fig. 1h**). Additional strain relief points were added along the Mesoscope using heat shrink wrap to keep wires together (repeat as needed for your designs, keep in mind more heat shrink also adds additional weight to the device).
8. **Magnets and fastening nut attachment:** Attach two disk magnets (B07C8ZZ2K9, Amazon) to the bottom surface of the bottom housing (**Supplementary Fig. 1i)**.

## SOFTWARE INSTALLATION

9. **Miniscope software installation:** Follow the instructions listed on the Miniscope page to install the DAQ software successfully on your computer. It will require installing the Miniscope software in various steps (http://miniscope.org/index.php/Software_and_Firmware_Setup).
10. **Microcontroller software installation:** Download the Arduino software (https://www.arduino.cc/en/main/software). You will additionally need to download the teensy (microcontroller) software (https://www.pjrc.com/teensy/td_download.html). Follow the steps on the Teensy page aforementioned to install the teensy software successfully and interface it with the Arduino library.

## WIRING

11. Connect the Mesoscope to the power circuit as shown in **Supplementary Figure 2a** and **b**. Follow the steps below to connect the power circuit controller to the computer and synchronize with the Miniscope CMOS sensor. **Ensure all ground wires from the DAQ and power supplies are connected to the common ground in the Mesoscope PCB**.
12. Solder two wires to the signal and ground pads (https://imgur.com/a/hgzua), of the CMOS sensor DAQ. These will connect to the microcontroller “trig-in” port (**Supplementary Figure 2b)**. This will act as the external trigger to start and stop synchronized videos.
13. Strip both ends of a piece of coax cable and separate into signal and ground wires. Solder a wire to each of the separated signal channels on one end, which will connect to the microcontroller “TTL out” port (**Supplementary Figure 2b)**. Solder the SMA to connector PCB on the other end of the stripped coax cable, which will be attached to the J5 port on the DAQ. This will allow you to collect TTL frame capture signals from the Miniscope DAQ.
14. Set up the two power supplies and ensure they are turned off during setup. Connect both power supply negative leads to the ground rail for each power supply (**Supplementary Figure 2b)**. Connect the positive cable from one power supply to the blue power terminal on the control board. Connect the positive cable from the other supply to the green power terminal on the control board. You should have a complete illumination power circuit now where the LEDs are powered by each power supply.

## TESTING LEDS

15. Now we can test the assembled circuit on any fluorescent object. Connect the Teensy and DAQ data cables to the USB 3.0 cables on the computer. Open the Arduino .ino file called “focusing LED.ino” and upload it to the Teensy. Open the Miniscope software to get a real-time view of the fluorescent molecules. Switch on the green LED power supply and you should see the green LED switch on (and remain on). The power supply should be at around 3V. Focus the Mesoscope to the desired FOV via the course focusing mechanism and tighten the set screw when a desired focus is achieved. Ensure to not overtighten the screw as it can break the set screw slot. Note green light is used because it can illuminate the cortex at a low power and therefore can be left on without being pulsed at these low powers. If you can see in-focus fluorescent material, the system is working.

## USING THE MESOSCOPE IN AN EXPERIMENT

16. With the Mesoscope focused and the system is wired as shown in **Supplementary Figure 2a**, run the ‘spontaneous pulsing.ino’ Arduino file on the Teensy. Ensure all the power supply wiring is correct (especially that the external power supply and signals are sharing the common ground rail with the Teensy). Open the .ino program and set desired parameters.
17. Set **Led_warm_up** to 120 seconds (LEDs are pulsed at moderate powers so they take time to warm up). This means the first 2 or so minutes of the video rendering are usually discarded since the LED dynamics have not stabilized.
18. Set the **experiment time per trial** as desired. The LEDs can usually be run for 6-8 minutes at a time before they start to heat up, so keep this in mind for the total time.
19. Set the **total number of trials**. The LEDs will warm up and the experiment time for one trial will be added to the first trial (i.e. if LEDs are 120sec to warm up and experiment time is 10 seconds, the first experiment trial will be finished at 130 sec. If there are more trials, i.e. 2, the second trial will occur between 130-140 seconds as the LEDs do not need to be warmed up again).
20. Set the blue LED pulse time (20 ms is recommended for 30 Hz acquisition) and the green LED pulse time (4 ms is recommended for 30 Hz CMOS acquisition). Note that the LED pulse time should be less than the duration of the specified framerate (i.e. if the frame rate is 60 Hz, 1 frame is captured every 16.66 ms. As the LEDs are pulsed on and off between frames, the LED pulse duration has to be less than 16.66 ms (usually 5-6 ms less than the frame rate works well).
21. Upload the .ino code with the set parameters to the Teensy
22. LED pulsing is synchronized to the frame capture of the Miniscope CMOS. Open the Miniscope software and connect the Miniscope CMOS. Set the desired frame rate (60hz usually drops a lot of frames so may not be recommended) and set the gain to a desired value (usually in the 50s). Tick the “trigger ext” box (by the record button); this will allow the microcontroller to trigger a video.
23. In Arduino, navigate to Tools→Serial Monitor. This will prompt the user (you) to enter the character ‘y’ to trigger the camera and begin recording the trial once the character command has been entered. Check the settings are correct and begin the trial. Usually the first trial is a test trial where you adjust the gain and LED power settings to ensure you are within the CMOS dynamic range (no over-saturation in the FOV).
24. Adjust the LED powers until you have sufficient illumination. The blue LEDs are usually run around 9-10V. The green LED is usually set to 4-6V. If the green LED is not required or desired, simply leave this power supply off during testing.

## Supplementary Note 2: Stability of LED illumination intensity

Fluctuations in the collected fluorescence signals could be attributed to changes in neural activity but could also be caused by fluctuations in the illumination power of the LEDs. We tested the stability of light power output delivered by the LEDs. Each LED is driven at a 30% duty cycle (20ms every odd frame lasting ∼33.3ms) when images are acquired at 30Hz. Each blue LED delivered mean light power of 7.67mW at 30% duty cycle. Continuous illumination at these powers for the durations tested (3-4 minutes) resulted in negligible changes in average blue light output ∼0.76% variation around the mean. The green LED delivered mean light power of 0.11mW at 6% duty cycle. Continuous illumination at these powers for the durations tested (3-4 minutes) resulted in negligible changes in average green light output ∼0.33% variation around the mean.

Additional tests conducted on the collected videos using the CMOS sensor were checked to quantify the cross-bleeding of the illumination light sources into each other’s frames. The emission signal for GCaMP is proportional to the blue excitation light power provided (independent of green light provided). As such, during the decay cycle of the blue LEDs, the amount of GCaMP that is excited also decreases very rapidly, and thus the cross-over of the blue excitation light into the green frames that can elicit an emission response is negligible. On the contrary however, the green reflectance illumination provided is calcium-independent in the sense that the cortex will have a reflectance response to any proportion of green illumination provided. The green illumination light can reflect back through the emission filter and onto the CMOS sensor as the green LEDs power down during alternate LED pulsing and potentially bleed into the blue frame signal. To quantify this response during realistic *in vivo* testing conducted on anesthetized mice, blue LEDs were pulsed for 6 minutes and the green LED was turned on at the 4-minute mark. The change in the mean blue intensity during this turn-on period was no more than 0.34% at 4ms pulses of the green LED illumination, thus demonstrating a reduced signal cross-over between the two illumination sources.

Pulsing the LEDs at high powers above their operating voltages can lead to additional light intensity noise and long-term damage of the LED themselves. To mitigate this issue, 1/8-inch copper tape was soldered to the back of the LED heat sink pads and allowed for long-term illumination experiments to be conducted (up to 10-minute experiments) with negligible LED damage and noise.

In summary, at a frame capture of 30 Hz, the LEDs were alternately pulsed with 20 ms pulses of the blue LEDs and 4 ms pulses of the green LED to maximize illumination intensity output while minimizing bleeding signal cross-over. An additional 2-minute LED warm-up period was allocated at the start of each test to eliminate non-linearities in LED illumination (**Supplementary Figure 2d**). Notably, it is not possible to fully correct for hemodynamics with reflectance measurements using a single wavelength^1^. However, GCaMP6 signal is much brighter than the intrinsic signal in the tissue. Further, improvements could be made with the development of brighter indicators, such as GCaMP7, for transgenic mice, or with the use of red-shifted activity reporters, where such corrections are not necessary. Both improvements could be integrated via relatively simple changes to the optical filters.

## REFERENCES

1. Chen, T.-W. et al. Ultrasensitive fluorescent proteins for imaging neuronal activity. Nature 499, 295–300 (2013).

2. Gong, Y. et al. High-speed recording of neural spikes in awake mice and flies with a fluorescent voltage sensor. Science (80-.). 350, 1361–1366 (2015).

3. Adam, Y. et al. Voltage imaging and optogenetics reveal behaviour-dependent changes in hippocampal dynamics. Nature (2019) doi: 10.1038/s41586-019-1166-7.

4. Madisen, L. et al. Transgenic mice for intersectional targeting of neural sensors and effectors with high specificity and performance. Neuron 85, 942–958 (2015).

5. Daigle, T. L. et al. A Suite of Transgenic Driver and Reporter Mouse Lines with Enhanced Brain-Cell-Type Targeting and Functionality. Cell (2018) doi: 10.1016/j.cell.2018.06.035.

6. Vanni, M. P. & Murphy, T. H. Mesoscale Transcranial Spontaneous Activity Mapping in GCaMP3 Transgenic Mice Reveals Extensive Reciprocal Connections between Areas of Somatomotor Cortex. J. Neurosci. (2014) doi: 10.1523/JNEUROSCI.1818-14.2014.

7. Vanni, M. P., Chan, A. W., Balbi, M., Silasi, G. & Murphy, T. H. Mesoscale mapping of mouse cortex reveals frequency-dependent cycling between distinct macroscale functional modules. J. Neurosci. (2017) doi: 10.1523/JNEUROSCI.3560-16.2017.

8. Borden, P. Y. et al. Genetically expressed voltage sensor ArcLight for imaging large scale cortical activity in the anesthetized and awake mouse. Neurophotonics (2017) doi: 10.1117/1.nph.4.3.031212.

9. Allen, W. E. et al. Global Representations of Goal-Directed Behavior in Distinct Cell Types of Mouse Neocortex. Neuron (2017) doi: 10.1016/j.neuron.2017.04.017.

10. Wekselblatt, J. B., Flister, E. D., Piscopo, D. M. & Niell, C. M. Large-scale imaging of cortical dynamics during sensory perception and behavior. J. Neurophysiol. (2016) doi: 10.1152/jn.01056.2015.

11. Makino, H. et al. Transformation of Cortex-wide Emergent Properties during Motor Learning. Neuron (2017) doi: 10.1016/j.neuron.2017.04.015.

12. Orsolic, I., Rio, M., Mrsic-Flogel, T. D. & Znamenskiy, P. Mesoscale cortical dynamics reflect the interaction of sensory evidence and temporal expectation during perceptual decision-making. bioRxiv (2019) doi: 10.1101/552026.

13. Musall, S., Kaufman, M. T., Juavinett, A. L., Gluf, S. & Churchland, A. K. Single-trial neural dynamics are dominated by richly varied movements. Nat. Neurosci. (2019) doi: 10.1038/s41593-019-0502-4.

14. Shimaoka, D., Steinmetz, N. A., Harris, K. D. & Carandini, M. The impact of bilateral ongoing activity on evoked responses in mouse cortex. Elife (2019) doi: 10.7554/eLife.43533.

15. Gilad, A. & Helmchen, F. Spatiotemporal refinement of signal flow through association cortex during learning. Nat. Commun. (2020) doi: 10.1038/s41467-020-15534-z.

16. Mohajerani, M. H. et al. Spontaneous cortical activity alternates between motifs defined by regional axonal projections. Nat. Neurosci. (2013) doi: 10.1038/nn.3499.

17. Ferezou, I. et al. Spatiotemporal Dynamics of Cortical Sensorimotor Integration in Behaving Mice. Neuron (2007) doi: 10.1016/j.neuron.2007.10.007.

18. Fink, A. J., Axel, R. & Schoonover, C. E. A virtual burrow assay for head-fixed mice measures habituation, discrimination, exploration and avoidance without training. Elife (2019) doi: 10.7554/eLife.45658.

19. Guo, Z. V. et al. Procedures for behavioral experiments in head-fixed mice. PLoS One (2014) doi: 10.1371/journal.pone.0088678.

20. Pinto, L. et al. Task-Dependent Changes in the Large-Scale Dynamics and Necessity of Cortical Regions. Neuron (2019) doi: 10.1016/j.neuron.2019.08.025.

21. Murphy, T. H. et al. High-throughput automated home-cage mesoscopic functional imaging of mouse cortex. Nat. Commun. (2016) doi: 10.1038/ncomms11611.

22. Bolaños, F. et al. Mesoscale cortical calcium imaging reveals widespread synchronized infraslow activity during social touch in mice. bioRxiv (2018) doi: 10.1101/430306.

23. Meyer, A. F., O’Keefe, J. & Poort, J. Two distinct types of eye-head coupling in freely moving mice. bioRxiv (2020) doi: 10.1101/2020.02.20.957712.

24. Juczewski, K., Koussa, J. A., Kesner, A. J., Lee, J. O. & Lovinger, D. M. Stress and behavioral correlates in the head-fixed method. bioRxiv 2020.02.24.963371 (2020) doi: 10.1101/2020.02.24.963371.

25. Aghajan, Z. M. et al. Impaired spatial selectivity and intact phase precession in two-dimensional virtual reality. Nat. Neurosci. (2015) doi: 10.1038/nn.3884.

26. Ghosh, K. K. et al. Miniaturized integration of a fluorescence microscope. Nat. Methods (2011) doi: 10.1038/nmeth.1694.

27. Cai, D. J. et al. A shared neural ensemble links distinct contextual memories encoded close in time. Nature 534, 115–118 (2016).

28. Zong, W. et al. Fast high-resolution miniature two-photon microscopy for brain imaging in freely behaving mice. Nat. Methods 14, 713 (2017).

29. Skocek, O. et al. High-speed volumetric imaging of neuronal activity in freely moving rodents. Nat. Methods (2018) doi: 10.1038/s41592-018-0008-0.

30. de Groot, A. et al. NINscope: a versatile miniscope for multi-region circuit investigations. bioRxiv (2019) doi: 10.1101/685909.

31. Scott, B. B. et al. Imaging Cortical Dynamics in GCaMP Transgenic Rats with a Head-Mounted Widefield Macroscope. Neuron (2018) doi: 10.1016/j.neuron.2018.09.050.

32. Namiki, S., Sakamoto, H., Iinuma, S., Iino, M. & Hirose, K. Optical glutamate sensor for spatiotemporal analysis of synaptic transmission. Eur. J. Neurosci. (2007) doi: 10.1111/j.1460-9568.2007.05511.x.

33. Marvin, J. S. et al. An optimized fluorescent probe for visualizing glutamate neurotransmission. Nat. Methods (2013) doi: 10.1038/nmeth.2333.

34. Dana, H. et al. Thy1-GCaMP6 transgenic mice for neuronal population imaging in vivo. PLoS One 9, (2014).

35. Ghanbari, L. et al. Cortex-wide neural interfacing via transparent polymer skulls. Nat. Commun. (2019) doi: 10.1038/s41467-019-09488-0.

36. Silasi, G., Xiao, D., Vanni, M. P., Chen, A. C. N. & Murphy, T. H. Intact skull chronic windows for mesoscopic wide-field imaging in awake mice. J. Neurosci. Methods 267, 141–149 (2016).

37. Ma, Y. et al. Resting-state hemodynamics are spatiotemporally coupled to synchronized and symmetric neural activity in excitatory neurons. Proc. Natl. Acad. Sci. U. S. A. (2016) doi: 10.1073/pnas.1525369113.

38. Valley, M. T. et al. Separation of hemodynamic signals from GCaMP fluorescence measured with wide-field imaging. J. Neurophysiol. (2020) doi: 10.1152/jn.00304.2019.

39. Ramos, A. Animal models of anxiety: do I need multiple tests? Trends Pharmacol. Sci. (2008) doi: 10.1016/j.tips.2008.07.005.

40. Dubbs, A., Guevara, J. & Yuste, R. moco: Fast motion correction for calcium imaging. Front. Neuroinform. (2016) doi: 10.3389/fninf.2016.00006.

41. Ghanbari, L. et al. Cortex-wide neural interfacing via transparent polymer skulls. Nat. Commun. 10, 1500 (2019).

42. Mathis, A. et al. DeepLabCut: markerless pose estimation of user-defined body parts with deep learning. Nat. Neurosci. (2018) doi: 10.1038/s41593-018-0209-y.

43. Dash, M. B., Douglas, C. L., Vyazovskiy, V. V., Cirelli, C. & Tononi, G. Long-term homeostasis of extracellular glutamate in the rat cerebral cortex across sleep and waking states. J. Neurosci. (2009) doi: 10.1523/JNEUROSCI.5486-08.2009.

44. Abadchi, J. K. et al. Spatiotemporal patterns of neocortical activity around hippocampal sharp-wave ripples. Elife (2020) doi: 10.7554/eLife.51972.

45. Patti, C. L. et al. Effects of sleep deprivation on memory in mice: Role of state-dependent learning. Sleep (2010) doi: 10.1093/sleep/33.12.1669.

46. Colavito, V. et al. Experimental sleep deprivation as a tool to test memory deficits in rodents. Front. Syst. Neurosci. (2013) doi: 10.3389/fnsys.2013.00106.

47. Xie, Y. et al. Resolution of high-frequency mesoscale intracortical maps using the genetically encoded glutamate sensor iGluSnFR. J. Neurosci. (2016) doi: 10.1523/JNEUROSCI.2744-15.2016.

48. Mohajerani, M. H., McVea, D. A., Fingas, M. & Murphy, T. H. Mirrored bilateral slow-wave cortical activity within local circuits revealed by fast bihemispheric voltage-sensitive dye imaging in anesthetized and awake mice. J. Neurosci. (2010) doi: 10.1523/JNEUROSCI.6437-09.2010.

49. Li, P. et al. Measuring sharp waves and oscillatory population activity with the genetically encoded calcium indicator GCaMP6f. Front. Cell. Neurosci. (2019) doi: 10.3389/fncel.2019.00274.

50. Dana, H. et al. High-performance calcium sensors for imaging activity in neuronal populations and microcompartments. Nat. Methods (2019) doi: 10.1038/s41592-019-0435-6.

51. Dawson, T. M., Golde, T. E. & Lagier-Tourenne, C. Animal models of neurodegenerative diseases. Nature Neuroscience (2018) doi: 10.1038/s41593-018-0236-8.

52. Juneau, J., Duret, G., Robinson, J. & Kemere, C. Enhanced Image Sensor Module for Head-Mounted Microscopes*. in Proceedings of the Annual International Conference of the IEEE Engineering in Medicine and Biology Society, EMBS (2018). doi: 10.1109/EMBC.2018.8512387.

53. de Groot, A. et al. Ninscope, a versatile miniscope for multi-region circuit investigations. Elife (2020) doi: 10.7554/eLife.49987.

54. Barbera, G., Liang, B., Zhang, L., Li, Y. & Lin, D. T. A wireless miniScope for deep brain imaging in freely moving mice. J. Neurosci. Methods (2019) doi: 10.1016/j.jneumeth.2019.05.008.

55. Ma, Y. Analysis of Resting-State Neurovascular Coupling and Locomotion-Associated Neural Dynamics Using Wide-Field Optical Mapping. ProQuest Dissertations and Theses (2018).

56. Lerner, T. N. et al. Intact-Brain Analyses Reveal Distinct Information Carried by SNc Dopamine Subcircuits. Cell (2015) doi: 10.1016/j.cell.2015.07.014.

57. Dana, H. et al. Sensitive red protein calcium indicators for imaging neural activity. Elife (2016) doi: 10.7554/elife.12727.

58. Donaldson, P. D., Ghanbari, L., Rynes, M. L., Kodandaramaiah, S. B. & Swisher, S. L. Inkjet-Printed Silver Electrode Array for in-vivo Electrocorticography. in 2019 9th International IEEE/EMBS Conference on Neural Engineering (NER) 774–777 (2019). doi: 10.1109/NER.2019.8717083.

59. Ghanbari, L. et al. Craniobot: A computer numerical controlled robot for cranial microsurgeries. Sci. Rep. 9, 1023 (2019).

60. Rynes, M. L. et al. Assembly and operation of an open-source, computer numerical controlled (CNC) robot for performing cranial microsurgical procedures. Nat. Protoc. (2020) doi: 10.1038/s41596-020-0318-4.

61. Pinheiro-da-Silva, J., Fernandes Silva, P., Nogueira, M. & Luchiari, A. ESM 1. (2016).

62. Schindelin, J. et al. Fiji: An open-source platform for biological-image analysis. Nature Methods (2012) doi: 10.1038/nmeth.2019.

63. Singh, S., Bermudez-Contreras, E., Nazari, M., Sutherland, R. J. & Mohajerani, M. H. Low-cost solution for rodent home-cage behaviour monitoring. PLoS One (2019) doi: 10.1371/journal.pone.0220751.

## REFERENCES

1. Ma, Y. et al. Resting-state hemodynamics are spatiotemporally coupled to synchronized and symmetric neural activity in excitatory neurons. Proc. Natl. Acad. Sci. U. S. A. (2016) doi: 10.1073/pnas.1525369113.

