## Supplementary figures and images for "Miniaturized head-mounted device for whole cortex mesoscale imaging in freely behaving mice"

### Mesoscope Cross Section 1.JPG

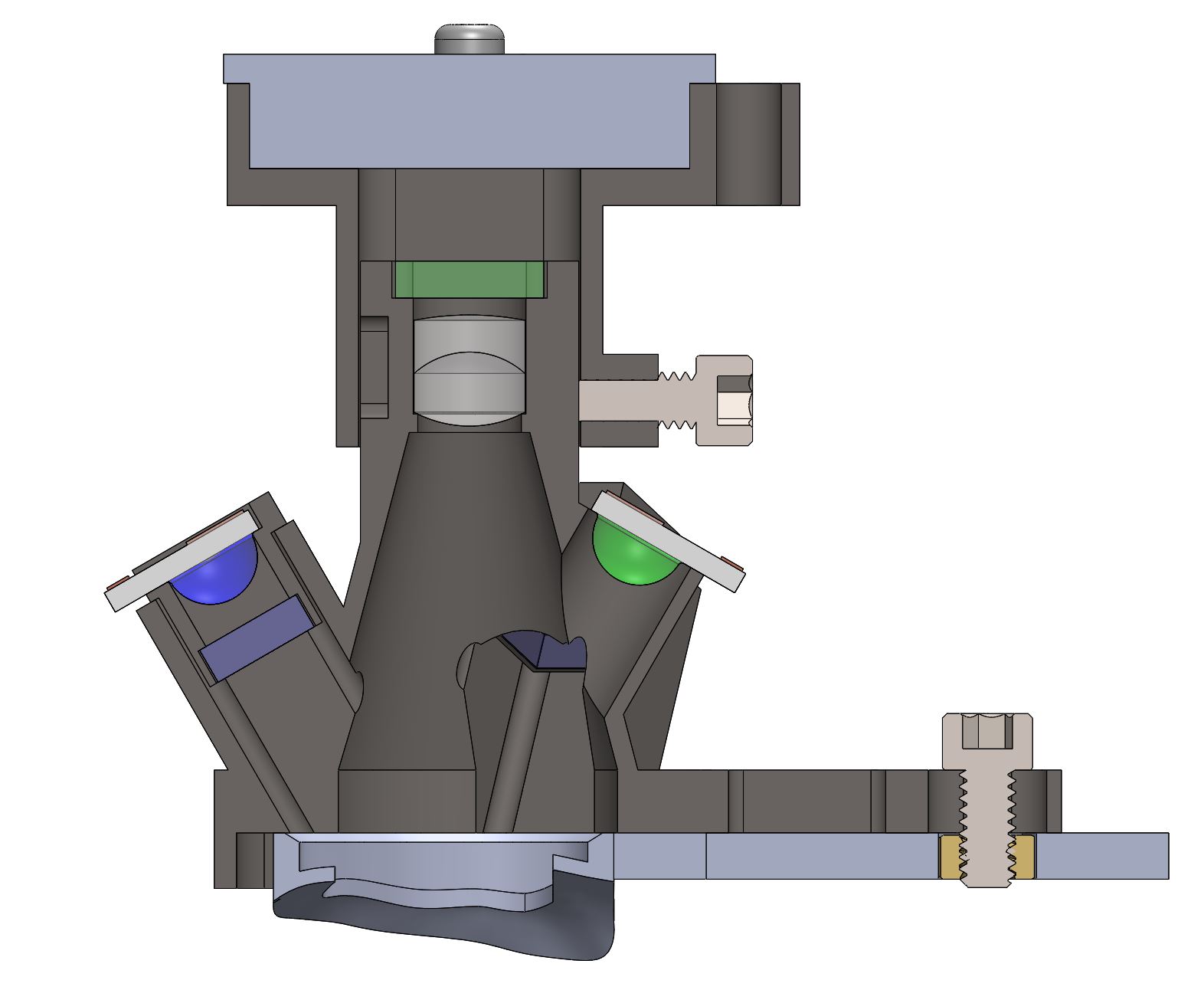

### Mesoscope Cross Section 2.JPG

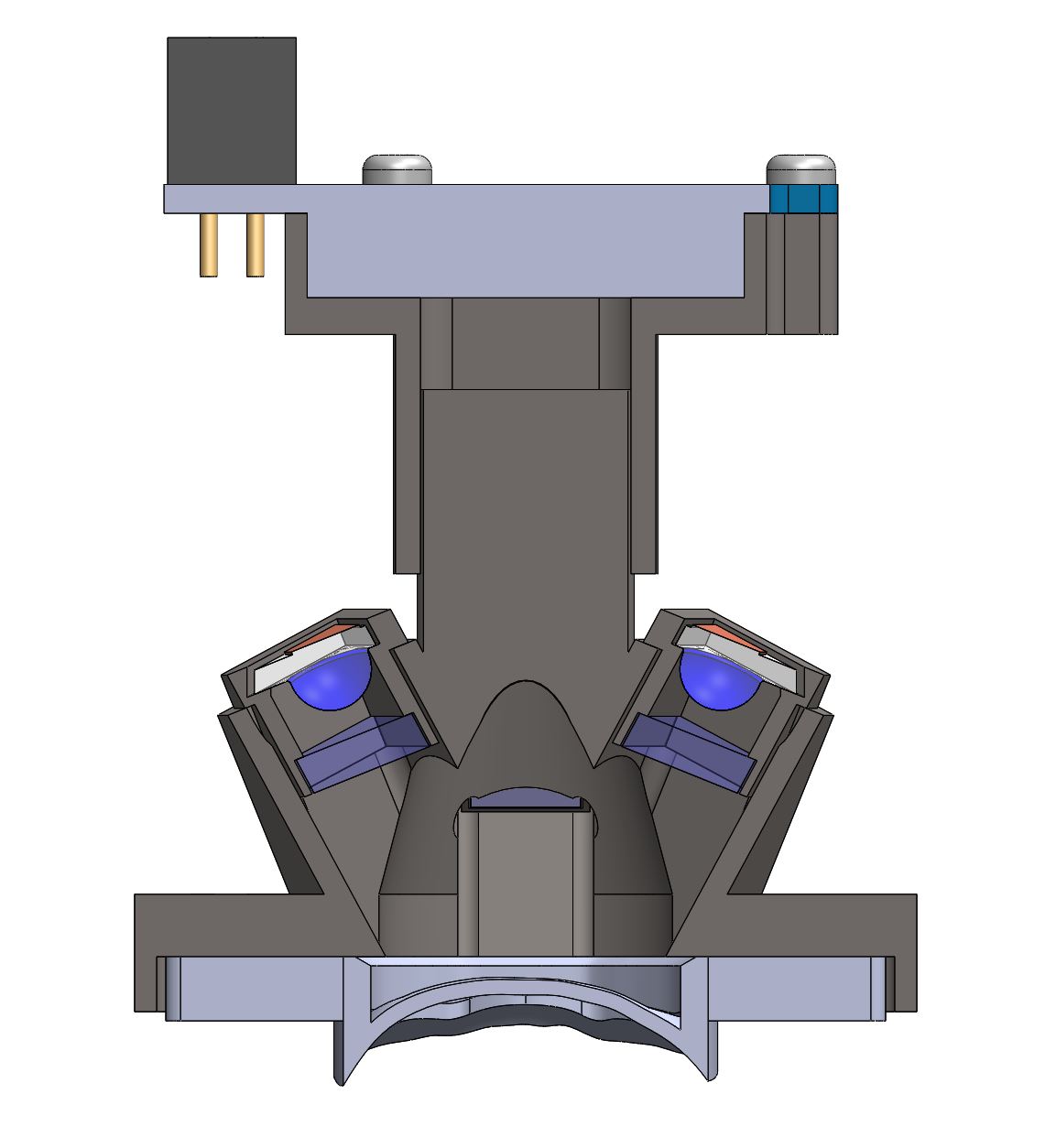

### Mesoscope Exploded View.JPG

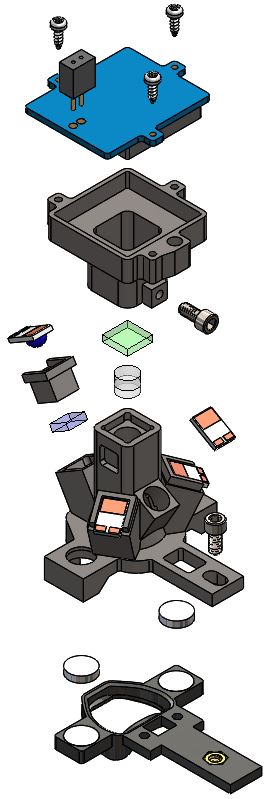
